# Myosin Heavy Chain Converter Domain Mutations Drive Early-Stage Changes in Extracellular Matrix Dynamics in Hypertrophic Cardiomyopathy

**DOI:** 10.1101/2022.04.05.487211

**Authors:** Jeanne Hsieh, Kelsie Becklin, Sophie Givens, Elizabeth R. Komosa, Juan E. Abrahante Lloréns, Forum Kamdar, Branden S. Moriarity, Beau R. Webber, Bhairab N. Singh, Brenda M. Ogle

## Abstract

More than 60% of hypertrophic cardiomyopathy (HCM)-causing mutations are found in the gene loci encoding cardiac myosin-associated proteins including myosin heavy chain (MHC) and myosin binding protein C (MyBP-C). Moreover, patients with more than one independent HCM mutation may be at increased risk for more severe disease expression and adverse outcomes. However detailed mechanistic understanding, especially at early stages of disease progression, is limited. To identify early-stage HCM triggers, we generated single (*MYH7 c*.*2167C>T* [R723C] with a known pathogenic significance in the MHC converter domain) and double (*MYH7 c*.*2167C>T* [R723C]; *MYH6 c*.*2173C>T* [R725C] with unknown significance) myosin gene mutations in human induced pluripotent stem cells (hiPSCs) using a base-editing strategy. Cardiomyocytes (CMs) derived from hiPSCs with either single or double mutation exhibited phenotypic characteristics consistent with later-stage HCM including hypertrophy, multinucleation, altered calcium handling, metabolism and arrhythmia. We then probed mutant CMs at time points prior to the detection of known HCM characteristics. We found *MYH7/MYH6* dual mutation dysregulated extracellular matrix (ECM) remodeling, altered integrin expression, and interrupted cell-ECM adhesion by limiting the formation of focal adhesions. These results point to a new phenotypic feature of early-stage HCM and reveal novel therapeutic avenues aimed to delay or prohibit disease onset.

## 1 Introduction

Hypertrophic cardiomyopathy (HCM) is the most common gene-linked cause of cardiac disease, affecting nearly 0.2% of the population globally (Semsarian et al., 2015). Clinical manifestations of HCM are highly variable, ranging from mild arrhythmias to sudden cardiac death (SCD), possibly due to single or multiple defects that give rise to myocyte disarray, fibrosis, hypertrophy, and heart failure (Marian and Braunwald, 2017). Among the many sarcomeric gene mutations associated with HCM, cardiac β-myosin heavy chain (β-MHC), myosin-binding protein C (MyBP-C), and cardiac Troponin T (cTnT) are the most prevalent (Burke et al., 2016; Marian and Braunwald, 2017). Some mutations result in early disease phenotypes, whereas others lead to late progressive HCM (Sedaghat-Hamedani et al., 2018). Whether variability in disease state is due to distinct molecular defects is largely unknown. While mutation type cannot currently be used to predict disease outcomes and guide management, patients with multiple independent sarcomere mutations have been identified to have more severe pathology including a higher prevalence of heart failure and SCD (Girolami et al., 2010; Kelly and Semsarian, 2009; Maron et al., 2012; Weissler-Snir et al., 2017). These findings highlight the need for comprehensive analysis using one or more HCM mutations. Here we focus our efforts on dual mutations in *MYH7* and *MYH6*.

*MYH7* encodes β-MHC and is composed of a globular head domain, a neck region, and a coiled-coil light meromyosin (LMM) domain (Homburger et al., 2016). The binding of Ca^2+^ exposes the myosin-binding sites on the actin filaments, leading to the formation of actin-myosin junctions and associated ability to glide along the actin base (Homburger et al., 2016). In the presence of ATP, these complexes are detached, and the ATPase activity of the head domain hydrolyzes ATP to initiate the next cycle of contraction (Homburger et al., 2016). A direct correlation between *MYH7* mutations, ATP utilization, and oxygen consumption has been established, however, whether these defects are associated only with head- or converter-domain mutation or both is not completely defined (Becker et al., 1997; Mosqueira et al., 2018). Although several HCM mutations are in the *MYH7* locus, significant clusters are found in the converter domain, also known as a “hot spot*”* of mutation (Homburger et al., 2016). Biophysical studies have shown that mutations in the converter region alter fiber stiffness, gliding velocity, and cross-bridge elasticity (Kawana et al., 2017; Kohler et al., 2002), thereby impacting the motor function of the myosin proteins; however, the functional impact of these mutations remains unclear.

*MYH6* gene encodes the other major MHC isoform, α-MHC. *MYH6* is expressed primarily in the atria in mature mammalian hearts and in the ventricle at embryonic stages and shares nearly 93% sequence homology with *MYH7* (McNally et al., 1989). Despite sequence and structural similarities between the α-MHC and β-MHC isoforms, α-MHC shows 150-300% higher contractile velocity and ∼70% higher affinity for the actin filaments than β-MHC proteins (Alpert et al., 2002; Locher et al., 2009). Like *MYH7* gene mutations, a number of HCM-causing mutations have been documented in the *MYH6* locus. For example, R795G results in late onset HCM (Niimura et al, 2002), and R721W showed 50% increased lifetime risk for sick sinus syndrome in carriers of this variant (Holm et al., 2011). Similar to *MYH7*, a mutation in the converter domain of *MYH6* is associated with congenital heart defects (Bjornsson et al., 2018). To our knowledge, phenotypic as well as mechanistic studies from such clinically-relevant *MYH6* mutations have not yet been performed.

hiPSC-derived cardiomyocytes (hiPSC-CMs) provide a pathophysiologically relevant approach to investigate *human* HCM and to design novel therapeutics (Lan et al., 2013; Mosqueira et al., 2018). Gene editing approaches further augment the utility of hiPSC-CMs to determine how certain mutations manifest the clinical disease phenotype. This approach also allows for studies of the mutation with an isogenic control to manage the genetic variability of individuals that may influence disease conditions (Bhagwan et al., 2020). Several studies have utilized gene-edited hiPSC-CMs to investigate the molecular details of HCM (Lan et al., 2013; Mosqueira et al., 2018; Yang et al., 2018). This approach is particularly valuable as available human tissue is largely limited to late-stage biopsy or cadaveric tissue. In addition, rodent models are of limited value for the current study since mice predominantly express *MYH6* in the heart as compared to *MYH7* in the human heart with significantly different cardiac electrophysiological properties relative to humans (Lowey et al., 2008). Recently, Mosqueira et al. demonstrated an HCM phenotype in *MYH7* R453C mutant hiPSC-CMs with sarcomeric disarray and arrhythmia (Mosqueira et al., 2018). Similarly, studies using *MYH7* E848G mutant hiPSC-CM showed disrupted interactions between β-MHC and MyBP-C with reduced contractility (Yang et al., 2018). These studies support the benefits of hiPSC-CMs to study HCM.

Here, we used hiPSC-CMs to evaluate the impact of converter domain mutations in the *MYH7* and *MYH6* locus on the pathogenicity of HCM and identified early ECM changes that precede later-stage physiologic defects associated with HCM. These studies are the first to link ECM dynamics with HCM onset and therefore provide a new avenue for drug discovery aimed at prevention or delay of disease symptoms.

## 2 Material and Methods

### 2.1 Gene Editing hiPSC Lines

#### 2.1.1 Guide RNA Design and Genomic DNA Analysis

The major isoform of *MYH7* for single-guide RNA (sgRNA) design was identified using NCBI (https://www.ncbi.nlm.nih.gov/) and the guide (gtatcgcatcctgaacccag) was chosen based on the on location of a PAM site placing the target C within the 5 base pair editing window (PMID: 27096365). The first passage after electroporation genomic DNA was isolated using QuickExtract™ (Lucigen), following manufacturer protocol. The target genomic locus was subsequently PCR amplified using AccuPrime Taq Hifi (ThermoFisher) (*MYH7* forward primer: cagtgacaaagccaggatca, reverse primer: ttggtgtggccaaacttgta; *MYH6* forward primer: gtgccccagagctcatagaa, reverse primer: ggcttctgcctcctaaactc) followed by Sanger sequencing of the PCR amplicon (ACGT). Sanger sequencing traces of base-edited samples were analyzed using EditR (z.umn.edu/editr) (PMID: 31021262). After clones were selected the top two off-target sites were sequenced following the same protocol as clonal selection (CPQ forward primer: agggaaccatggccataaatgt, reverse primer: gcctccctggggaagtgata; MYO5b forward primer: gccgagcctcagaagtgtag, reverse primer: ccccctcccagtagaactca).

#### 2.1.2 hiPSC Electroporation and Clonal Isolation

Two hours prior to electroporation, hiPSCs were treated with 10µM ROCK inhibitor (Y27632 dihydrochloride, Tocris). Cells were then passaged and 250K cells were then spun down in PBS. Cells were resuspended in P3 Primary Cell Nucleofector™ Solution containing Supplement 1 (Lonza, Switzerland) and 1 µg of chemically modified sgRNA (Synthego, Menlo Park, CA) and 1.5 µg codon optimized BE4 mRNA (TriLink Biotechnologies, San Diego, CA) and placed in the 20µl cuvette provided and CB-150 was the Amaxa protocol used. Cells were given 10µM ROCKi for 48 hours post electroporation followed by normal culturing conditions. For hiPSC single cell isolation ∼300 single cell suspended iPSC were cultured in a 10cm dish with 10µM ROCKi for 48 hours. For the next ten days cells were given daily media changes and then colonies were isolated using a 10µl pipette to scrape half of the colony for isolated culture and half the colony for genomic DNA isolation.

### 2.2 Cardiac Differentiation

Isogenic control hiPSC-CMs and mutant hiPSC-CMs were obtained via the fully defined direct cardiac differentiation by modulating Wnt/b-catenin signaling. Briefly, hiPSCs were cultured on Matrigel-coated 6-well plates in mTeSR1 medium for 5 days to reach 80-90% confluency. To harvest singularized cells, the culture medium was removed, and the wells were washed one time with sterile DPBS, then added 1 mL of room-temperature Accutase into each well. The plates were transferred into the incubator to allow Accutase to react at 37°C for 8 min. 0.5 mL of mTeSR1 medium was added into each well to stop Accutase reactions. The singularized cells were resuspended in mTeSR1 with 5 μM ROCK inhibitor (Y27632), seeded with the density of 5 × 10^5^ cells/well on Matrigel-coated 24-well plates and maintained at 37°C, 5% CO2 incubator for 24 hours. After 24 hours, on Day -1, 1 mL of fresh mTeSR1 medium was added to each well. On Day 0, the start of cardiac differentiation, 1 mL of RPMI/B-27 without insulin (RPMI minus) medium with 8 μM GSK3-β inhibitor (CHIR99021) was added into each well for 24 hours in the 37°C, 5% CO2 incubator to induce mesoderm. On Day 1, CHIR99021 medium was replaced with fresh RPMI/B27 minus medium. On Day 3, 0.5 mL of exhausted medium from each well was collected and mixed 1:1 with fresh RPMI/B27 minus insulin medium. Wnt inhibitor (IWP2) was added into the mixed medium to achieve the concentration of 5 μM. Each well was incubated with IWP2 for 48 hours to induce cardiac differentiation. On Day 5, fresh RPMI/B27 minus insulin was added into each well. On Day 7 and every 3 days thereafter, fresh maintenance medium, RPMI/B-27 medium, was added to each well. Spontaneous twitching was observed on Day 8, and robust spontaneous contraction occurred by Day 12. To purify the cardiomyocyte population, lactate enrichment was performed on Day 15-18 via adding 1 mL/well of DMEM (without glucose) with 4 mM sodium L-lactate every 2 days. On Day 19 and every 3 days thereafter, cells were maintained in RPMI/B27 medium.

### 2.3 Immunohistochemical Staining

Cultured hiPSC-CMs were fixed with 4% paraformaldehyde (PFA) for 15 min and then washed and stored in PBS at 4°C. To perform immunostaining, cultured hiPSC-CMs were permeabilized by 0.2% Triton X-100 at room temperature for 1 hr, washed with PBS 3 times, and then added blocking buffer (2.5g non-fat dry milk in 50 mL of 0.2% Triton X-100) to incubate at room temperature for 2 hrs. Washed with PBS 3 times before adding primary antibody (mouse anti-cTnT, 1:200 dilution in BGST buffer) for overnight incubation at 4°C. Washed with 0.2% Tween 20 for 2 times and PBS for 2 times before adding secondary antibody (Alexa Fluor 647 goat anti-mouse IgG, 1:500 dilution in BGST buffer), plates were covered by aluminum foil and incubated at room temperature for 1.5 hrs. Washed with 0.2% Tween 20 for 2 times and PBS for 2 times before staining for nuclei with 0.1μg/mL of DAPI dye for 15 min at room temperature. Washed with PBS 3 times to remove the excess dye and stored in PBS at 4°C before imaging. For co-staining of pFAK and cTnT, we followed the protocol above with slight modifications. Briefly, after fixation, we permeabilized the cardiomyocytes using 0.2% Triton X-100 in PBS (0.2% PBST) for 10 minutes followed by washing with 0.1% PBST three times. The cells were blocked using 5% BSA/0.1% PBST for 1hr and incubated with primary antibodies, cTnT-1:300; pFAK-1:1000 in the blocking solution in a humified chamber overnight. Subsequently, the cells were washed 4 times with 0.1% PBST followed by incubation with secondary antibodies (1:500) for 1 h at room temperature. Cells were washed 4 times and stained for nuclei using DAPI (1:1000), and images using Leica fluorescence microscope.

### 2.4 ADP/ATP Assay

ADP/ATP assay was performed as per manufacturer’s instructions using ADP/ATP Ratio Assay Kit (Bioluminescent) (Abcam, ab65313). Isogenic control and mutant hiPSCs were differentiated to cardiomyocytes as described above and single-cell suspension was obtained at each time point. A total of 50000 cells from each condition for each time point were re-suspended in the 250 uL of nucleotide releasing buffer for 5 min at room temperature. A total of 50 μL of the cell lysate was added to the reaction mix and incubated for 2 min before measuring the ATP levels. To measure the ADP levels, 10 μL of ADP converting enzyme was added to the reaction mix and fluorescence was measured after 2 min of incubation using the bioluminescence plate reader.

### 2.5 RNA Isolation and qPCR Analysis

Total RNA were isolated from undifferentiated and differentiated isogenic control and R723C hiPSCs using the RNeasy kit (Qiagen) according to the manufacturer’s protocol. Briefly, washed the cells once with sterile PBS and lysed in RLT-lysis buffer for specific time points. RNA isolation was performed as per the protocol, followed by on-column DNA digestion to remove any traces of DNA as per the instructions. For cDNA synthesis, 100-1000 ng of total RNA was used and synthesized using the SuperScript IV VILO kit (Thermo Fisher Scientific) according to the manufacturer’s protocol. Quantitative PCR (qPCR) was performed using gene-specific oligos and the SYBR-green method. The list of oligos used is provided in Table S3.

### 2.6 Bulk-RNA Sequencing and Analysis

The total RNA was isolated as described above. For the bulk-RNAseq analysis, 1 μg of total RNA was submitted to Novogene (https://en.novogene.com/) for sequencing and data analysis. Downstream analysis was performed using a combination of programs including STAR, HTseq, Cufflink, and our wrapped scripts. Alignments were parsed using Tophat program and differential expressions were determined in R using the package DESeq2/edgeR. GO and KEGG enrichment was implemented by the ClusterProfiler. Gene fusion and difference of alternative splicing events were detected by Star-fusion and rMATS software. Reference genome and gene model annotation files were downloaded from the genome website browser (NCBI/UCSC/Ensembl) directly. Indexes of the reference genome were built using STAR and paired-end clean reads were aligned to the reference genome using STAR (v2.5). STAR used the method of Maximal Mappable Prefix (MMP) which can generate a precise mapping result for junction reads. HTSeq v0.6.1 was used to count the read numbers mapped of each gene. And then FPKM of each gene was calculated based on the length of the gene and the reads count mapped to this gene. FPKM, Reads Per Kilobase of exon model per Million mapped. reads, considers the effect of sequencing depth and gene length for the reads count at the same time, and is currently the most used method for estimating gene expression levels. Differential expression analysis between two conditions/groups (two biological replicates per condition) was performed using the DESeq2 R package (2_1.6.3). DESeq2 provides statistical routines for determining differential expression in digital gene expression data using a model based on the negative binomial distribution. The resulting P-values were adjusted using the Benjamini and Hochberg’s approach for controlling the False Discovery Rate (FDR). Genes with an adjusted P-value <0.05 found by DESeq2 were assigned as differentially expressed. To identify the correlation between differences, we clustered different samples using expression level FPKM to see the correlation using the hierarchical clustering distance method with the function of the heatmap, SOM (Selforganization mapping), and kmeans using silhouette coefficient to adapt the optimal classification with default parameter in R.

### 2.7 Statistical Analysis

All experiments were repeated at least two times and the values presented are mean ± standard error of the mean (SEM) from the replicates of at least two independent experiments. Statistical significance was determined using the Student’s t-test when comparing 2 groups and ONE-WAY ANOVA with multiple comparisons when comparing more than 2 groups. A p-value < 0.05 was considered a significant change and was highlighted in each panel by an asterisk.

## 3 Results

### 3.1 *MYH7* R723C mutation can result in an early pathogenic HCM phenotype

Mutations in the *MYH7* locus result in severe HCM phenotypes in human patients (Burke et al., 2016) especially those in the converter domain, a “hot spot” for HCM mutations (Table S1). We identified a total of 14 patients with a single mutation of *MYH7* R723C using database searches (Table S2). There was gene penetration in 64% (9/14) patients. The mean age of the gene penetrant patients was 29 ± 13 years and maximal wall thickness was 24 ± 10 mm (range 11-38 mm). Three patients were described to have symptoms which included heart failure. Patient 1 was symptomatic at age 12 and underwent cardiac transplant at the age of 16. Patient 1’s father was the proband and had the same mutation and end-stage heart failure, but further clinical information was not available to include him in the study. Tesson et al. described an Italian family with 11 members with a *MYH7* R723C mutation in which the proband was incidentally identified on routine screening. In this family, patients 8 and 9 were monozygotic twins and gene penetrance was undetermined for patient 9 given ECG changes but normal indexed wall thickness. Above clinical observations indicate the *MYH7* R723C mutation can result in early pathologic HCM to asymptomatic incomplete penetrance. Additional clinical data on this mutation coupled with *in vitro* mechanistic studies will be important to understand gene penetrance, symptoms, and survival. Further, no study has explored the possible compensatory impact of wild-type a-MHC protein in these patients given the structural similarity of b-MHC to a-MHC. In the current study, we investigate the mechanisms that govern initial onset of HCM in the presence of pathogenic *MYH7* mutation and another mutation in the corresponding position of the *MYH6* locus with unknown clinical significance.

### 3.2 Generation and characterization of isogenic control and MHC mutant hiPSC lines

Multiple reports have shown the pathological symptoms of c.2167C>G (R723G) and c.2167C>T (R723C) mutations in *MYH7* gene locus leading to hypertrophic cardiomyopathy in human patients (Homburger et al., 2016; Kraft et al., 2016). Interestingly, while the R723G mutation is well-described using biochemical, structural, and gene-editing approaches (Homburger et al., 2016; Kawana et al., 2017; Kraft et al., 2016), the impact of R723C mutation remains unclear, and the functional and mechanistic details are not yet defined. The arginine residue at the 723 location is highly conserved across multiple species (figure S1A, B) and mutation in this region has been shown to alter mRNA secondary structure of *MYH7* (Rose et al., 2020). To investigate the earlier stage MHC mutation-associated HCM pathology and to account for the effect of wild-type α-MHC expressed by immature human induced pluripotent stem cell-derived cardiomyocytes (hiPSC-CMs) in 2D culture system, we utilized CRISPR/Cas9 base editing to generate a homozygous *c*.*2167C>T* mutation in the *MYH7* locus (figure 1A, figure S2A-C) with an analogous heterozygous *c*.*2173C>T* mutation in the *MYH6* locus in a hiPSC line reprogrammed from cardiac fibroblasts of a healthy female donor and modified to overexpress cyclin D2 to enhance differentiation yield (Gaudelli et al., 2017; Webber et al., 2019). We identified and expanded a clonal population of mutated cells with the targeted gene mutations of cytosine (C) to thymidine (T) at *c*.*2167C>T* and *c*.*2173C>T* positions. Sequencing showed a single chromatogram peak at the targeted positions, with no off-target effect (figure 1B). Initially, we evaluated the pluripotency of the gene-edited lines using the *MYH7* R723C and *MYH6* R725C (*MYH7/MYH6*) dual mutant hiPSCs and the corresponding isogenic control hiPSCs. Immunostaining using OCT4 antibody showed no difference between control and mutant lines (figure 1C). Further, qPCR analysis demonstrated similar expression of *OCT4* transcripts in the control and mutant lines (figure 1D). These results indicate that the stemness of mutant hiPSCs was not affected by the mutation or base editing procedure.

**Figure 1.**
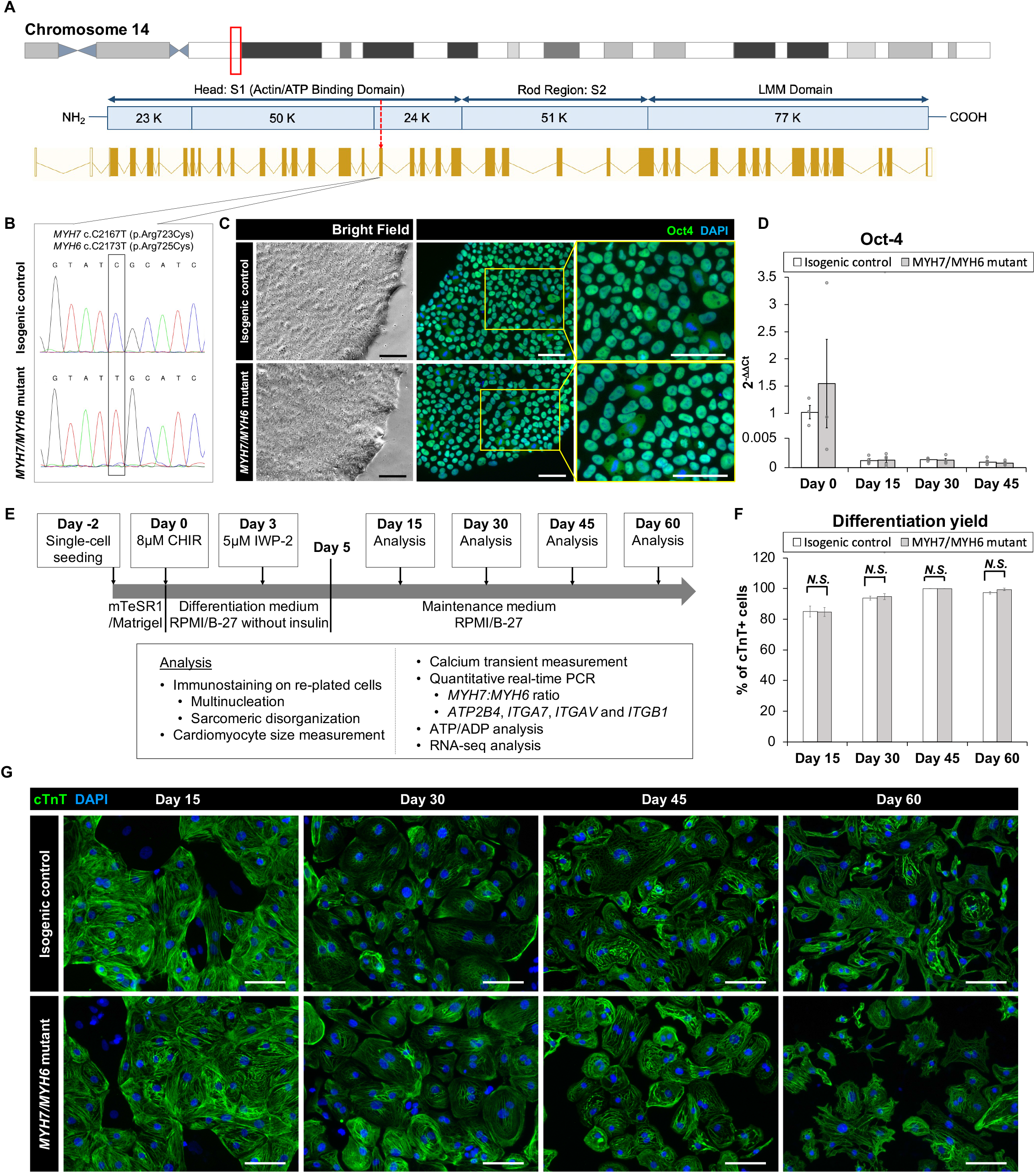
Base editing and characterization of the *MYH7/MYH6* mutant hiPSCs. **(A)** Schematic of *MYH7* locus as shown in the red box on chromosome 14 and c.2167 location as pointed by the red arrow for base editing. **(B)** Chromatogram of *MYH7* mutant locus in hiPSCs showing c.C2167T (p.Arg723Cys, p.R723C) mutation and its isogenic parental genotype. **(C)** Bright field images and immunostaining of Oct-4 using the isogenic control and *MYH7/MYH6* mutant hiPSCs. DAPI labels the nuclei (blue). **(D)** qPCR analysis for Oct-4 expression level using RNA isolated from the isogenic control and *MYH7/MYH6* mutant hiPSCs. **(E)** Cardiac differentiation timeline and analysis regime. **(F)** Quantitative analysis of the differentiation efficiency based on the cTnT staining in the control and *MYH7/MYH6* mutant cardiomyocytes. Note that mutation did not result in any significant difference in the cardiomyocyte differentiation efficiency. **(G)** Cardiac Troponin T (cTnT) and DAPI-stained hiPSC-CMs on Day 15, 30, 45 and 60 after the start of differentiation. Data represents mean ± SEM. (Scale Bar = 100 μm) (*N*.*S*.: not significant).

To assess the cardiomyocyte differentiation potential of the *MYH7/MYH6* mutant, single *MYH7* R723C (*MYH7*) mutant and the isogenic control lines, we differentiated the lines using a small-molecule protocol (Lian et al., 2012) and performed an array of analyses as schematized in figure 1E. Spontaneous beating was observed between Day 8 and Day 12 of differentiation in both control and mutant lines. As expected, the *OCT4* transcripts were greatly reduced at Day 15 relative to Day 0 and remained low throughout the differentiation (figure 1D, Day 15-45). Next, we assessed the efficiency of cardiomyocyte differentiation using immunofluorescence staining for cardiac troponin T (cTnT) up to Day 60 of differentiation (figure 1F, G). Quantification of cTnT+ cells relative to DAPI-labeled nuclei revealed no significant difference in the differentiation efficiency between control and *MYH7/MYH6* mutant hiPSC-CMs (figure 1F). As indicated by the percentage of cTnT+ cells, both mutant and the control lines exhibited more than 85% differentiation efficiency before lactate purification (Day 15) and more than 99% differentiation efficiency after lactate purification (Day 30, 45, and 60). There was no difference on the percentage of cTnT+ cells among the hiPSC-CMs of the *MYH7* mutant, *MYH7/MYH6* mutants, and the isogenic controls (figure S3A). These results indicate that the *MYH7* and *MYH7/MYH6* mutations did not affect cardiomyocyte differentiation potential.

### 3.3 *MYH7/MYH6* and *MYH7* mutant hiPSC-CMs recapitulate HCM cellular phenotypes

Several studies have utilized patient-derived or gene-edited hiPSC-CMs to model HCM phenotypes *in vitro* with supportive as well as variable findings (Bhagwan et al., 2020; Kraft et al., 2016; Lan et al., 2013; Yang et al., 2018). These differences may be dependent upon the type of mutation and location (Bhagwan et al., 2020). To evaluate the impact of *MYH7/MYH6* and *MYH7* mutations on HCM phenotypes, we carried out a series of studies comparing the mutant hiPSC-CMs to the corresponding isogenic control hiPSC-CMs. Initially, we undertook analyses of cellular-scale features over time. To evaluate cellular hypertrophy, we analyzed the singularized cells prior to attachment to well plates and found that both *MYH7/MYH6* mutant cells and *MYH7* mutant cells were significantly larger in size than the isogenic control cells (figure S3B, S3C). We next counted the number of nuclei per cell and found a significant increase in multinucleated cells in the mutant hiPSC-CMs (figure S3D). At Day 15 of differentiation, mutant hiPSC-CMs had slightly lower number of multinucleated cells than the isogenic control hiPSC-CMs, while at Day 45 of differentiation, both *MYH7/MYH6* and *MYH7* mutant hiPSC-CMs had statistically significant higher number of multinucleated cells than the isogenic lines.

With little difference in cellular scale features among the two *MYH7/MYH6* mutant lines and one *MYH7* mutant line, we focused remaining comparisons on one isogenic control line and one *MYH7/MYH6* mutant line for a more detailed study of HCM phenotype progression over 4 different time points (i.e., Day 15, 30, 45 and 60 of differentiation). In this way we accounted for possible functional compensation of *MYH6* expressed at early developmental time points and correspondingly muted HCM phenotype in the *MYH7* single mutant. We found that while both control and *MYH7/MYH6* mutant hiPSC-CMs increased cardiomyocyte size over time, *MYH7/MYH6* mutant hiPSC-CMs were significantly larger than the control hiPSC-CMs at all time points (figure 2A, B). This hypertrophic phenotype was further verified using FACS analysis of the singularized cardiomyocytes from control and *MYH7/MYH6* mutant lines between Day 15-45 (figure S4A, B). Regarding multinucleation, quantitative analysis revealed that *MYH7/MYH6* mutant hiPSC-CMs had significantly more multinucleated cells than the control hiPSC-CMs from Day 30 of differentiation, and the difference was most pronounced at Day 45 with a two-fold increase in the *MYH7/MYH6* mutant condition (27 ± 5%) relative to control (12 ± 3%) (figure 2E).

**Figure 2.**
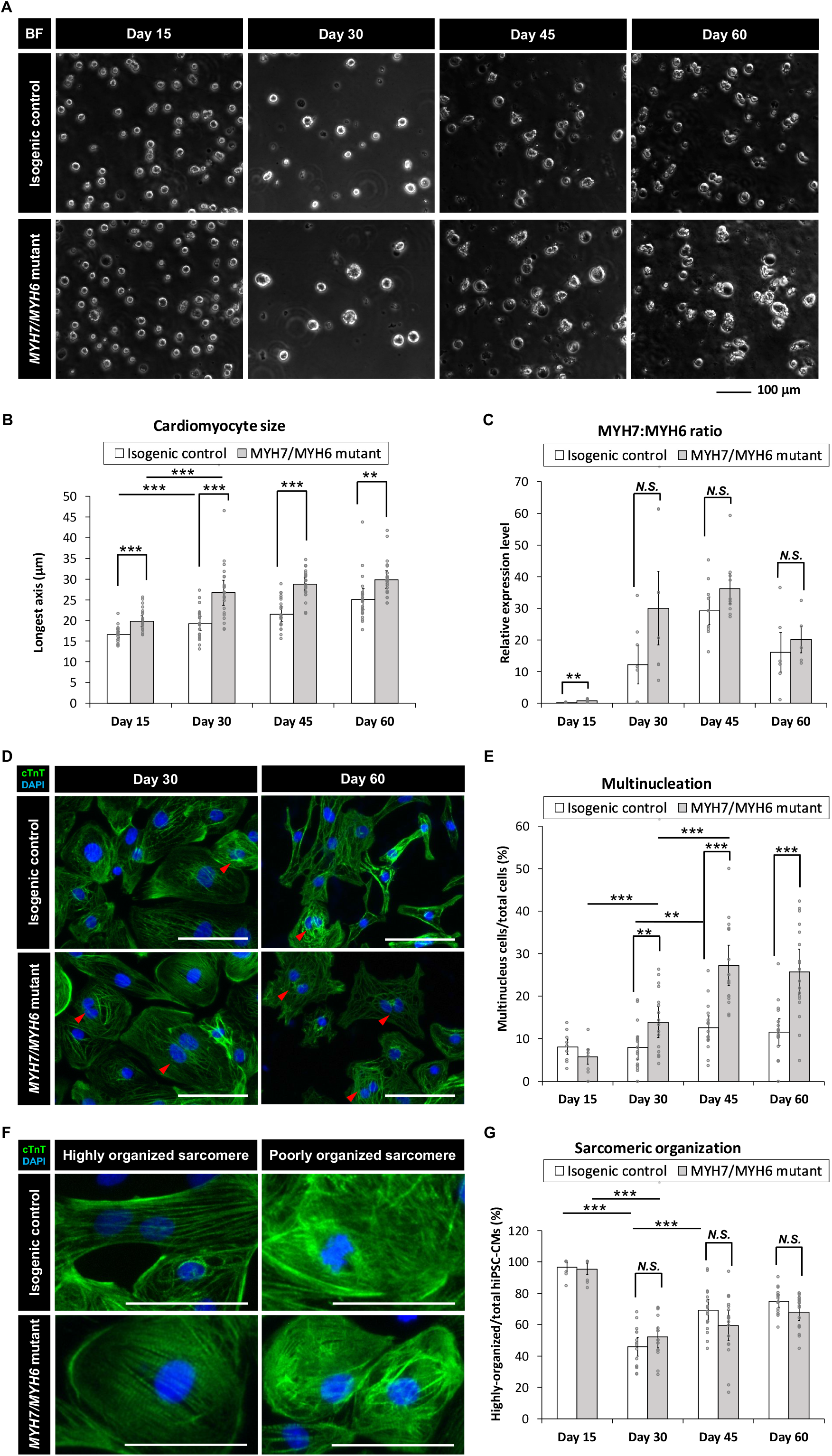
*MYH7/MYH6* mutation recapitulates the hypertrophic cardiomyopathy phenotype in the hiPSC-CMs. **(A, B)** Bright-field images **(A)** and quantification (longest axis) **(B)** of the re-plated control and mutant cardiomyocytes prior to their attachment to the substrate to analyze their size at Day 15-60 of differentiation. Notice the hypertrophy of the mutant cells from Day 15-60. (Scale Bar = 100 μm). Data represents mean ± SEM. (\*\**P*<0.01; \*\*\**P*<0.005) **(C)** *MYH7*:*MYH6* ratio obtained from qPCR analysis. Data represents mean ± SEM. (\*\**P*<0.01; *N*.*S*.: not significant) **(D)** Representative images of cTnT-immunostained and DAPI-stained hiPSC-CMs to visualize nucleation in the differentiated cardiomyocytes using the isogenic control and *MYH7/MYH6* mutant lines. Red arrows indicate multi-nucleated cells. Left images show Day 30 re-plated hiPSC-CMs and Right images show Day 60 re-plated hiPSC-CMs. (Scale Bar = 100 μm). **(E)** Bar graph shows that quantification of the percentage of hiPSC-CMs with more than one nucleus. Note significant increase in the multinucleated cells in the mutant conditions relative to control cells. Data represents mean ± SEM. (\*\**P*<0.01; \*\*\**P*<0.005). **(F)** High magnification fluorescence images of cTnT and DAPI-stained hiPSC-CMs to analyze the sarcomeric organization in the control and R723C mutant cardiomyocytes. (Scale Bar = 50 μm). **(G)** Bar graph shows percentage of hiPSC-CMs with highly organized sarcomere. Data represents mean ± SEM. (\*\*\**P*<0.005; *N*.*S*.: not significant).

We then assessed the ratio of *MYH7*:*MYH6* transcripts using qPCR and found an increase in the mutant cardiomyocytes relative to the control isogenic lines (figure 2C). A similar increase in the *MYH7*:*MYH6* ratio has been documented in hypertrophic patients (Lowes et al., 1997; Mosqueira et al., 2018). Further, our analysis revealed that the increase in *MYH7*:*MYH6* ratio was mainly caused by a reduced level of expression of *MYH6* (less energy efficient) relative to the control, without much change in *MYH7* (more energy efficient) transcripts. Another HCM phenotype reported with *MYH7* mutations is sarcomere disorganization and myofibrillar disarray; however, we did not find any difference in the percentage of cells with irregular and unorganized sarcomeric structures in the hiPSC-CMs from control and mutant cells at any of the time points examined (figure 2F, G). We also found a gradual but unexpected decline in the overall cell density in the mutant lines relative to the control lines after the lactate purification (figure S5). Aside from altered cell density and lack of unorganized sarcomeres, these data reflect the progressive nature of HCM phenotypes in the MHC-mutant hiPSC-CMs, as seen in HCM patients.

### 3.4 *MYH7/MYH6* and *MYH7* mutations result in functional impairment of hiPSC-CMs

Having established HCM cellular phenotypes, we next studied the impact of *MYH7/MYH6* mutation on cardiomyocyte function. We found that the spontaneous beating rates were significantly lower in the mutant cardiomyocytes as compared to the control hiPSC-CMs between Day 15-60 of differentiation (figure 3A, B and video S1). Comparable results of lower spontaneous beat rates were observed in the *MYH7* mutant hiPSC-CMs (figure S3E). The beat rates of the control hiPSC-CMs linearly increased over time, whereas the beat rates of mutant hiPSC-CMs were not significantly changed between Day 15-45 with a slight decline at Day 60 of differentiation. Further, the maximal spontaneous beat rates of control hiPSC-CMs were significantly higher than the mutant hiPSC-CMs at Day 45 of differentiation (figure 3B, S3E). Notably, we found statistically more events of irregular beating (arrhythmia) in the mutant hiPSC-CMs at Day 30 and Day 45 of differentiation (figure 3A, C-D). These results indicate contractility defects associated with *MYH7/MYH6* mutation in cardiomyocytes. Since contractility is dependent upon the Ca^2+^ ion concentration, we next performed calcium transient measurements using Fluo-4AM fluorescent dye. Our analysis showed a significantly higher calcium signal in the mutant hiPSC-CMs relative to the control hiPSC-CMs at all time points examined (figure 3E), indicative of increased recruitment of Ca^2+^ ions in the mutant cardiomyocytes. Further, similar to the trend of beat rate over time, calcium signal reached maximal value at Day 45 and then declined at Day 60 in the mutant hiPSC-CMs (figure 3B and 3E). These results establish the importance of the converter domain of *MYH7/MYH6* in the regulation of CM functionality. Also, our result indicates that mutations in the converter domain, such as *MYH7* R723C mutation, affects cross-bridge interactions as well as Ca^2+^ ion dependency.

**Figure 3.**
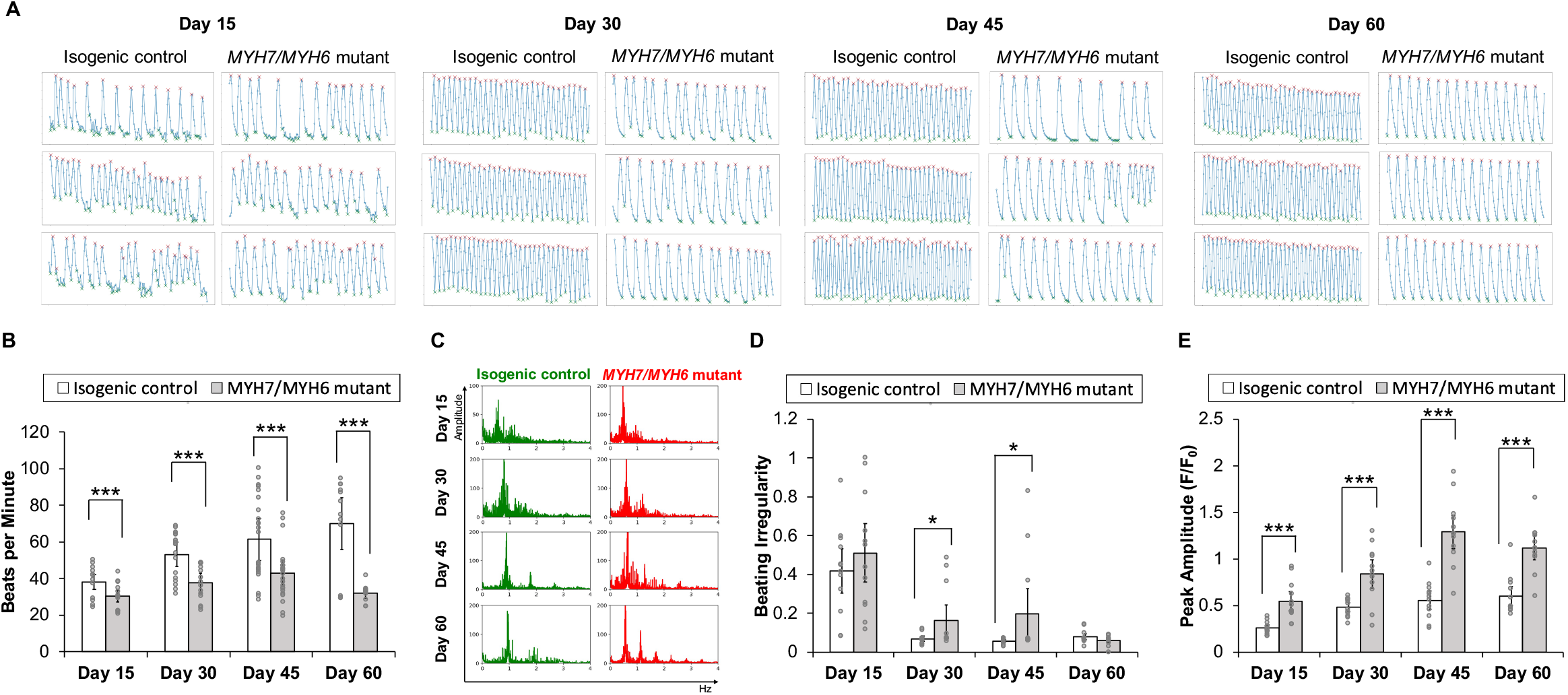
Functional assessment of the *MYH7/MYH6* mutant cardiomyocytes. **(A)** Beat rate measured from calcium transient analysis using the isogenic control and *MYH7/MYH6* mutant cardiomyocytes at Day 15-60 of differentiation. Note the significant reduction in beats per minute in the mutant cardiomyocytes relative to control. Data represents mean ± SEM; \*\*\**P*<0.005. **(B)** Beat rate in frequency domain for both isogenic control and *MYH7/MYH6* mutant cardiomyocytes at Day 15-60 of differentiation. **(C)** Quantitative result for irregular beating in the isogenic control and *MYH7/MYH6* mutant cardiomyocytes at Day 15-60 of differentiation. The level of beating irregularity was defined as the average standard deviation for all beat intervals over twelve calcium transient measurements for each cell line at each time point. Data represents mean ± SEM; \**P*<0.05. **(D)** Peak amplitude of calcium signal measured from calcium transient analysis. Data represents mean ± SEM; \*\*\**P*<0.005.

### 3.5 *MYH7/MYH6* mutation leads to alterations in mitochondrial activity and ATP/ADP turn-over

To investigate *MYH7/MYH6* mutation-associated molecular details at the early stage of HCM development, we performed bulk RNA-seq analysis at Day 15 of differentiation using RNA isolated from the isogenic control and mutant hiPSC-CMs. We performed pair-end sequencing with >30 million reads per sample with clean read counts of more than 96% for each sample (figure S6). Downstream analysis was performed using a combination of programs including STAR, HTseq, Cufflink, and alignments were parsed using Tophat and differential expressions were determined through DESeq2/edgeR R package (3.16.5). The resulting heatmap comparing HCM mutant and control hiPSC-CMs is shown in figure 4A. From RNA-seq analysis, we identified a total of 3,317 differentially expressed genes (DEGs); of these 49% were up-regulated genes and 51% were down-regulated genes in the mutant relative to the control (figure 4A-C). Gene ontology analysis using DisGeNET as well as Disease Ontology (DO) database resources revealed robust enrichment of transcripts involved in hypertrophic cardiomyopathy together with other myopathy-related transcripts (figure 4D, E). Next, our KEGG analysis showed that transcripts up-regulated in the mutant cardiomyocytes were related to hypertrophic cardiomyopathy (*ACTC1, MYL2, MYL3*), Ca^2+^-dynamics regulatory pathway (*ATP2B4, CACNA1C, CAMK2B, PLN*), as well as cardiac-muscle contraction transcripts (*ACTC1, MYH7, TNNI3, TNNT2*) (figure 4F-H). These studies supported our initial cellular and functional assay in the recapitulation of HCM phenotype in *MYH7/MYH6* mutant cardiomyocytes.

**Figure 4.**
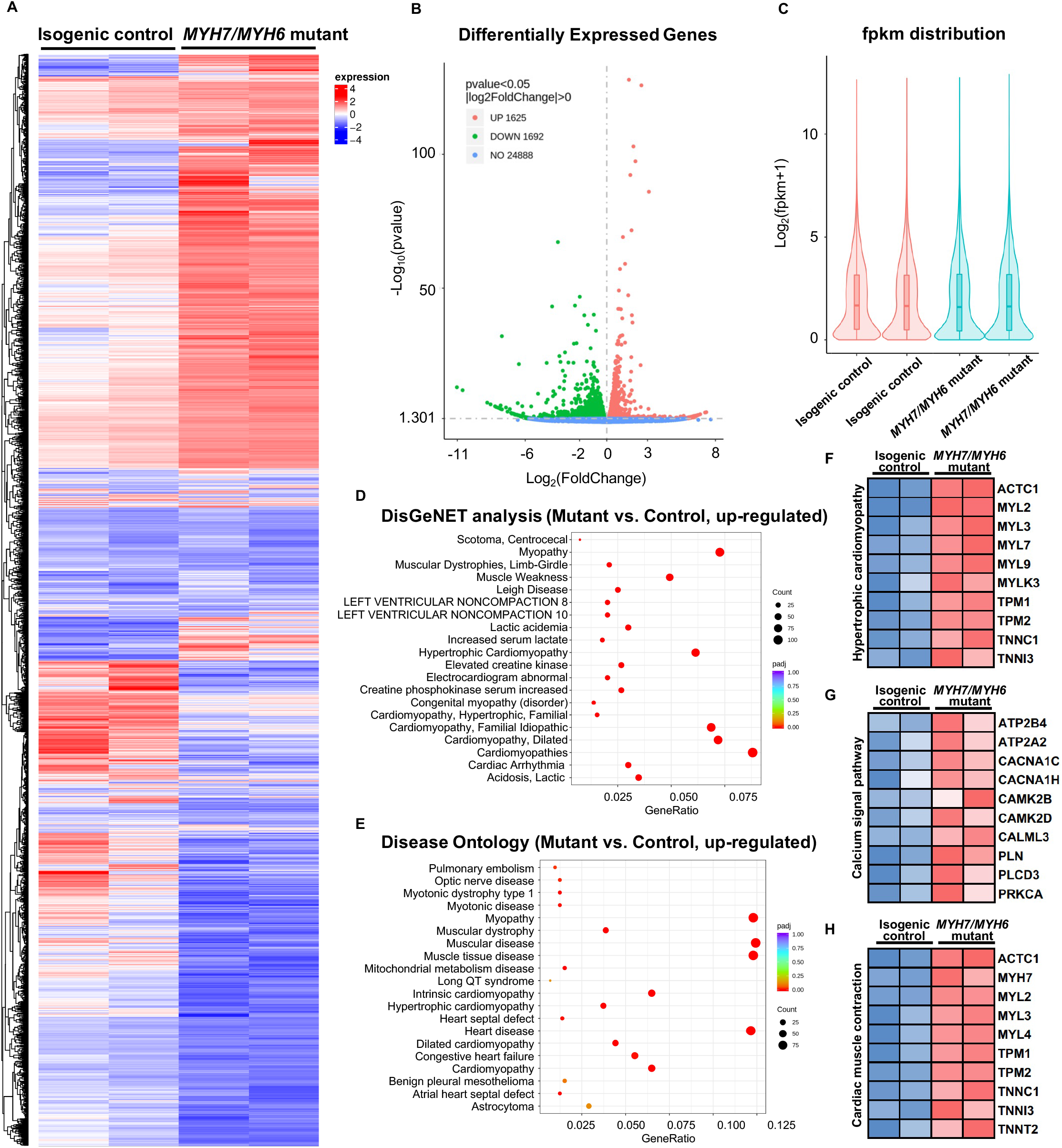
Bulk RNAseq analysis of *MYH7/MYH6* dual mutation-induced hypertrophic cardiomyopathy in hiPSC-CMs to evaluate the molecular mechanism. **(A)** Heatmap for comparing HCM mutant and control hiPSC-CMs showing the dysregulated transcripts in the mutant cardiomyocytes relative to control. **(B)** Volcano plot showed a total of 3,317 differentially expressed genes (DEGs) with the mutation. (P<0.05, |log_2_fold change|>0) **(C)** fpkm distribution showing similar log counts between the control and mutant lines. **(D)** Top 20 up-regulated gene-disease associations by DisGeNet analysis. **(E)** Top 20 up-regulated diseases annotated by disease ontology (DO) analysis. Size of the dot indicate the number of gene associated with the given disease and red color indicate the significance of this association. **(F)** Heatmap for 10 up-regulated DEGs that associated with hypertrophic cardiomyopathy. **(G)** Heatmap for 10 up-regulated DEGs that associated with calcium signal pathway. **(H)** Heatmap for 10 up-regulated DEGs that associated with cardiac muscle contraction.

It has been shown that different HCM mutations result in inefficient sarcomeric ATP utilization, energy depletion, and increased ATPase activity of the β-MHC protein (Lan et al., 2013; Mosqueira et al., 2018). To evaluate whether a mutation in the converter domain altered any of these properties, we analyzed the RNAseq datasets for mitochondrial and metabolic pathways. Interestingly, we found robust enrichment of transcripts related to the TCA cycle, respiratory electron-transport pathway, pyruvate metabolism, mitochondrial biogenesis, and complex I biogenesis in the mutant hiPSC-CMs relative to the isogenic control hiPSC-CMs (figure 5A). Additional analysis revealed up-regulation of several mitochondrial transcripts including *MT-ND1-6, MT-CYB, MT-ATP6* in the mutant hiPSC-CMs (figure 5B). To further support these findings, we performed an ATP/ADP bioluminescence assay using cardiomyocytes from control and mutant lines at various time points. We did not observe any difference in the ADP:ATP ratio until Day 30 of differentiation relative to control hiPSC-CMs. However, we found a robust and sharp increase at Day 45, indicating a surge in the need for ATP utilization at later time points in the mutant hiPSC-CMs as compared to control (figure 5C). Next, we evaluated the expression of *ATP2B4* (an ATPase, Ca^2+^/Mg^2+^-dependent enzyme) using RNA isolated from control and mutant lines at Day 15 and Day 45 of differentiation. Similar to ADP:ATP ratio, we found a robust enrichment of *ATP2B4* mRNA at Day 45 of differentiation in the mutant cardiomyocytes relative to control cardiomyocytes (figure 5D). Overall, these data further substantiate the progressive nature of the *MYH7/MYH6* dual mutation-mediated HCM phenotypes.

**Figure 5.**
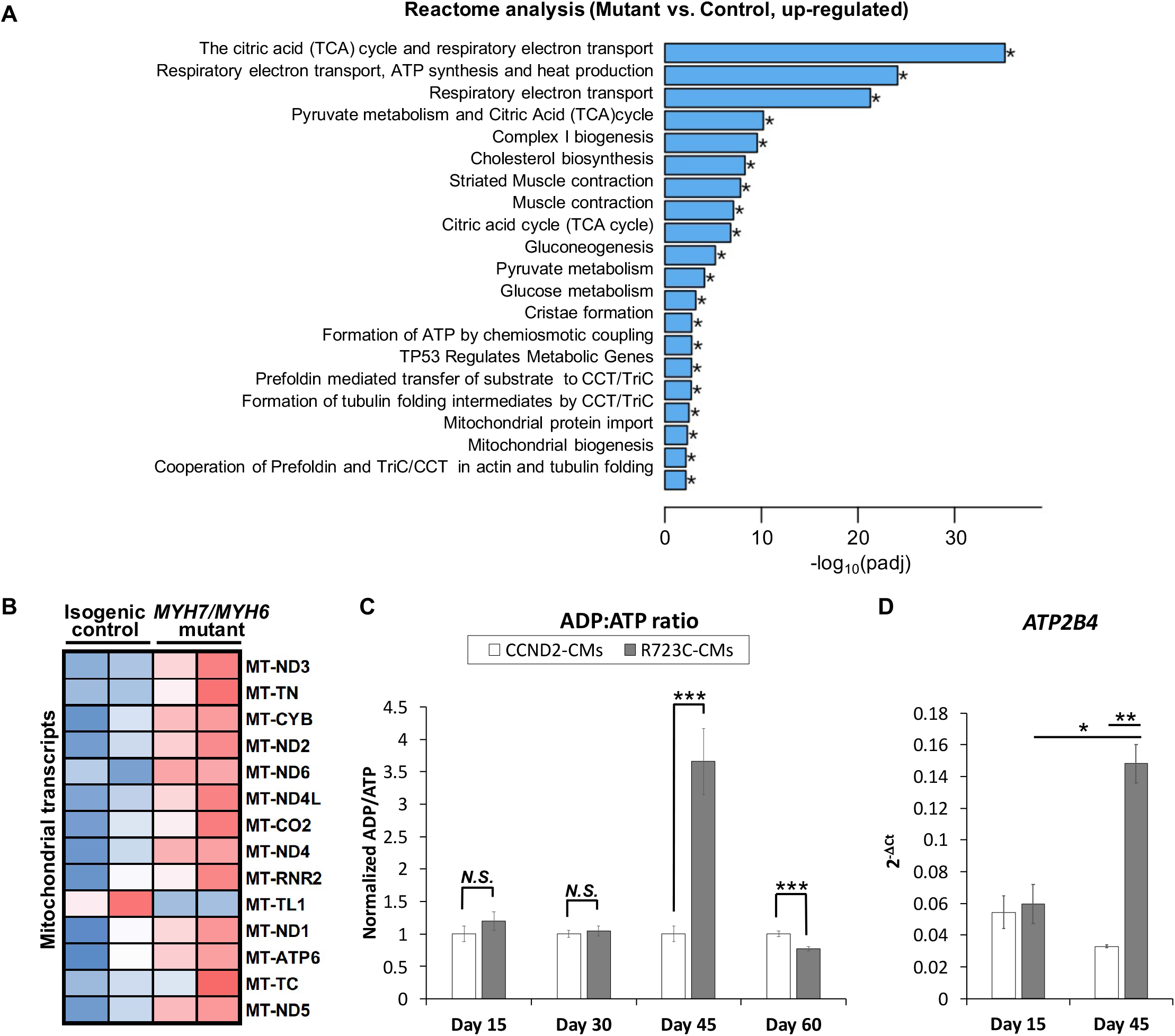
*MYH7/MYH6* mutation results in dysregulation of the metabolic pathways and ADP/ATP dynamics. **(A)** Bar graph showing top 20 significantly altered metabolic pathways in the mutant cardiomyocytes relative to control cardiomyocytes; \**P*<0.05. **(B)** Heatmap of the DEGs mitochondrial transcripts in the mutant cardiomyocytes relative to control cardiomyocytes. **(C)** ADP:ATP ratio using the lysate from control and mutant cardiomyocytes. Notice the sharp peak at Day 45 of differentiation in the mutant cells relative to control cells. Data represents mean ± SEM; \*\*\**P*<0.005; *N*.*S*.: not significant. **(D)** qPCR analysis for ATP2B4 transcripts using the RNA isolated from control and mutant cardiomyocytes at Day 15 and Day 45 of differentiation. Data represents mean ± SEM; \**P*<0.05; \*\**P*<0.01.

### 3.6 *MYH7/MYH6* mutation leads to changes in ECM dynamics

Our analysis of the bulk RNAseq datasets revealed several down-regulated transcripts in the mutant line including those related to pathways involved in fibrosis, focal adhesion, and adherent junctions (Varnava et al., 2000). Among several pathways, our Reactome analysis revealed that transcripts related to ECM were most significantly down-regulated in the mutant hiPSC-CMs (figure 6A, B), suggesting for the first time that ECM dynamics play a critical role in the development of the HCM phenotype. Our heatmap analysis showed dysregulation of several ECM transcripts such as down-regulation of integrin subunits *ITGA2, ITGA4, ITGA8, ITGA9, ITGB8*, and up-regulation of *ITGA7, ITGAV, ITGB1, JUN* (figure 6B). Next, our KEGG analysis revealed that genes required for the ECM-receptor interactions were dysregulated in the *MYH7/MYH6* mutant hiPSC-CMs (figure 6C, D). Because ECM-receptor interaction was the most significantly down-regulated pathway and because loss of cardiomyocytes (i.e., low cell density) was observed in mutant hiPSC-CMs at later time points after the differentiation (figure 6C, D and figure S5), we investigated whether the interaction of the integrins with downstream intracellular signaling pathways and their ability to attach to the Matrigel matrix is altered in the mutant cardiomyocytes. The binding of ECM to integrins induces multiprotein clustering at focal adhesions and mediates intracellular signaling through focal adhesion kinase (FAK) (Guan, 1997). Upon activation, FAK undergoes autophosphorylation (pFAK) and forms a complex with Src and other cellular proteins to trigger downstream signaling through its kinase activity or scaffolding function (Guan, 1997, 2010). To evaluate this, we performed co-staining using cTnT and pFAK antibodies in the control and mutant hiPSC-CMs. Interestingly, our data revealed activated FAK signaling (pFAK staining), punctate staining near the membrane regions of the control hiPSC-CMs (figure 6E). Notably, this activation was significantly reduced in the mutant hiPSC-CMs with little staining in the membrane regions (figure 6E). Next, to investigate ECM-cell dynamics, among the differentially expressed integrin genes, we selected *ITGB1, ITGAV*, and *ITGA2* for flow cytometric analysis to monitor protein level over time. Our data showed up-regulation of *ITGB1* at Day 30 and Day 45 of differentiation and down-regulation of *ITGA2* at Day 30 without a significant change in *ITGAV* levels (figure S7A-C). Further, our qPCR analysis revealed higher expression of *ITGAV* and *ITGB1* transcripts using RNA isolated from isogenic control and mutant cardiomyocytes at Day 45 (figure 6F). These studies support the hypothesis that the *MYH7/MYH6* mutation disrupts cell-ECM interactions, leading to a stressed state and perturbed function of cardiomyocytes.

**Figure 6.**
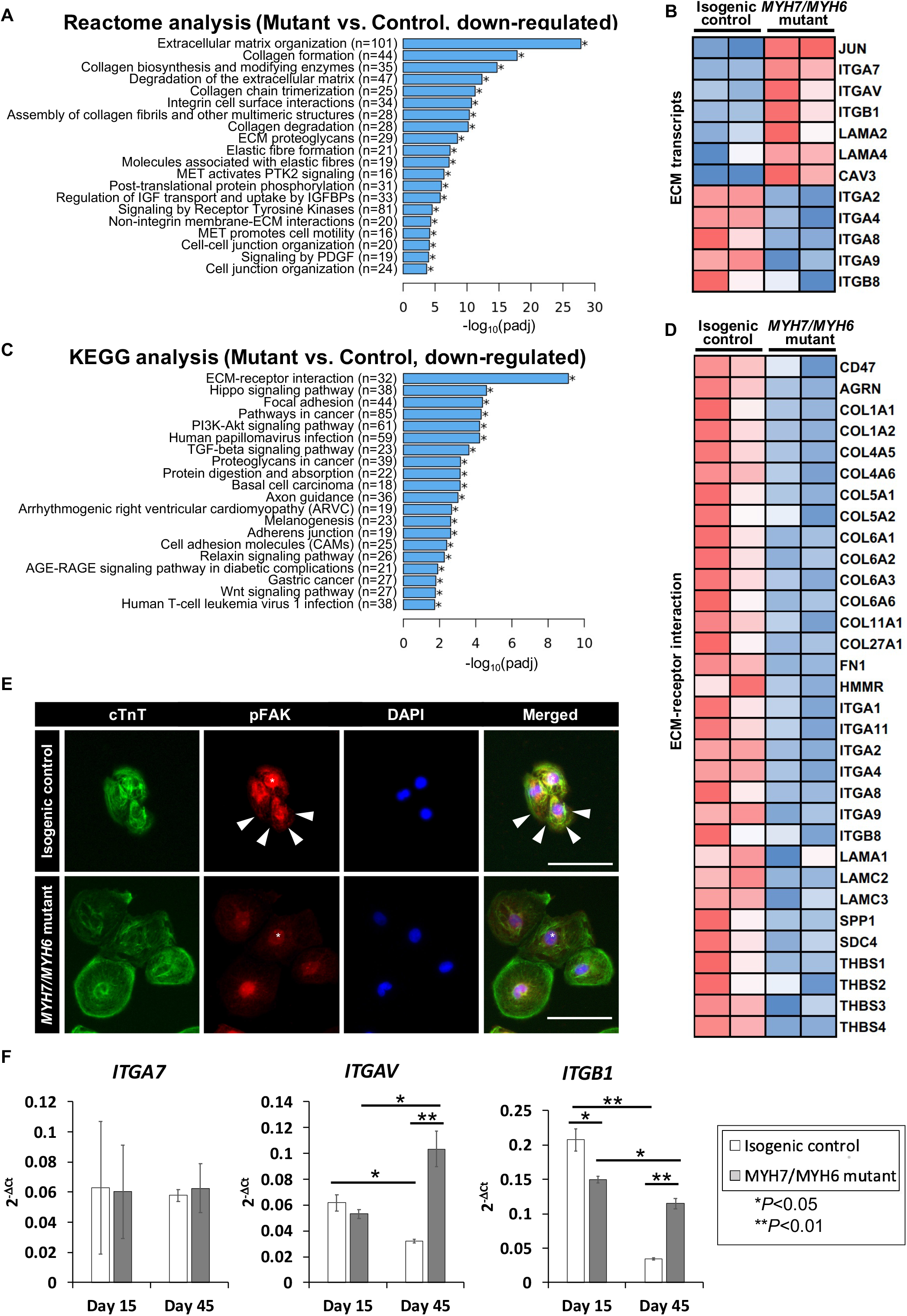
*MYH7/MYH6* mutation drives altered ECM dynamics and ECM-receptor interaction. **(A)** Reactome analysis showing the top 20 annotated down-regulated pathways in the mutant as compared to control lines. Note that the most significantly dysregulated were the pathways associated with ECM organization. **(B)** Heatmap showing the dysregulated transcripts involving in ECM. **(C)** Top 20 annotated pathways determined by KEGG analysis with down-regulated DEG list. Note that ECM-receptor interaction transcripts were most significantly altered in the mutant cells relative to control cells. **(D)** Heatmap for up-regulated DEGs that associated with ECM-receptor interaction. **(E)** Representative images of cTnT and pFAK-immunostained hiPSC-CMs using isogenic control and mutant cells. White arrows indicate activated FAK signal at the vicinity of the cell-substrate attachment site. DAPI labels the nuclei (blue). (Scale Bar = 100 μm) **(F)** qPCR analysis for *ITGA7, ITGAV* and *ITGB1* transcripts using RNA isolated from control and mutant cardiomyocytes at Day 15 and Day 45 of differentiation. Data represents mean ± SEM; \**P*<0.05; \*\**P*<0.01.

## 4 Discussion

Because HCM is mostly due to sarcomeric protein mutations, novel approaches for investigating HCM genotype-phenotype correlations and molecular mechanisms are essential to identify potential therapeutic targets. In this study, we reviewed the clinical phenotype of the *MYH7* R723C mutation, engineered *MYH7* R723C and *MYH6* R725C mutant hiPSCs using base-editing method, and compared HCM phenotypes in the mutant and corresponding isogenic control hiPSC-CMs. The use of hiPSC-CMs as a model system allowed us to study *early-stage* cell and molecular alterations that give rise to well-described later-stage phenotypes and clinical symptoms of HCM (Sedaghat-Hamedani et al., 2018).

The most significant early-stage molecular alterations detected in *MYH7/MYH6* mutant lines were in ECM remodeling and ECM-receptor interactions. RNAseq, qPCR, and immunostaining data revealed perturbed ECM synthesis and degradation, expression of integrins, and activation of FAK. Integrins are surface receptors and are required for cell-cell interactions as well as cell-matrix interactions (Guan, 1997). Binding of ECM to the integrin receptors is essential for proper intracellular signaling and activation of multi-protein cascades (Guan, 2010). Integrins *ITGAV, ITGA7, ITGB1* were up-regulated in the mutant hiPSC-CMs, while many ECM genes and others integrins were down-regulated. The possibility that these early changes initiate the pathogenesis of the HCM phenotype was further substantiated by the fact that mutant cardiomyocytes became more sparsely dispersed on the plate at later time points perhaps as a function of altered cell-ECM adhesion (figure S5). In further support, a study using *MYH7* R403Q mutation documented the impact of a diseased matrix in prolonged contraction and poor relaxations in engineered heart tissue (EHT) and in a swine model (Sewanan et al., 2019). We depict this new insight regarding early changes in cell-ECM communication as a mediator HCM disease in figure 7. Future studies will be needed to determine whether these outcomes are substantiated in the 3D environment and with mechanical stimulation.

**Figure 7.**
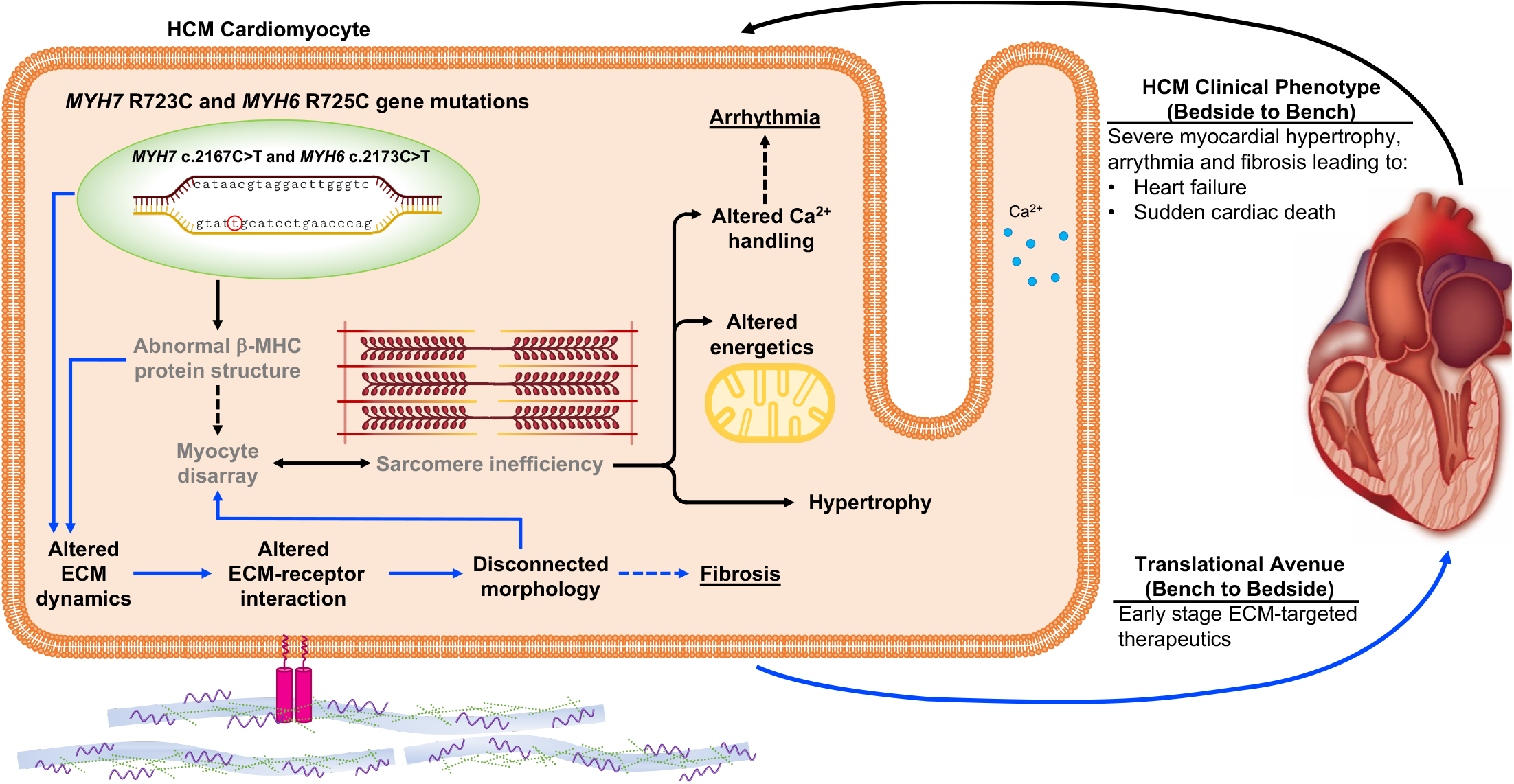
Schematic depicting the impact of *MYH7/MYH6* mutation in mediating hypertrophic cardiomyopathy. Our data indicate that *MYH7/MYH6* mutation results in hypertrophy, dysregulation of metabolic as well as ECM dynamics. Solid black arrows and bold text represent HCM phenotypes that have been reported for other *MYH7* mutations, grey text represents previously reported features of *MYH7* mutations. Solid blue arrows represent the mutation associated ECM dynamics which is novel to the field and could provide targets to develop drug therapies potent in the early stages of HCM.

We propose drug therapeutics that target these ECM-receptor interactions at the early stages of human development as a new means to prevent or delay the onset of HCM.

We also defined changes in the metabolic pathways and corresponding mitochondrial transcripts in the mutant cardiomyocytes relative to the control. While the observation that metabolic pathways are altered with HCM is not new, we add further detail. In particular, we found an aberrant and robust peak in the ADP/ATP dynamics in the mutant hiPSC-CMs at Day 45 of differentiation. However, the defects in beat rate and Ca^2+^ amplitude were significantly different starting from Day 15 and plateauing by Day 45. Based on these results, we postulate that in the mutant hiPSC-CMs, there is increased utilization of ATP by the myosin head to continue the next cycles of contraction. These events lead to the accumulation of ADP in the cells, and increased ADP/ATP ratio. Next, we propose that the decline in the ADP/ATP ratio at Day 60 could be due to a secondary mechanism such as apoptosis or fibrosis in the mutant cardiomyocytes. Indeed, we found several fibrotic transcripts as well as cell death machinery transcripts. While these results provide strong evidence for functional defects and ATP utilization in the *MYH7/MYH6* dual mutation-associated HCM, we also recognize that other factors such as fiber stiffness (Kohler et al., 2002) and contraction velocity (Kawana et al., 2017) might also contribute to this phenotype and will be the focus of future studies.

We also provide evidence for increased mitochondrial load and changes in metabolic profiling in the mutant hiPSC-CMs. Whether this is due to the *MYH7/MYH6* mutations in the converter domain of MHC with increased fiber stiffness, the requirement of higher energy, or imperfect interactions of actin-myosin bridging is not clear from this study. Further biochemical assays will be needed to answer these questions. It has been shown that other *MYH7* mutations, R723G as well as R719W, resulted in a 15% increase in gliding velocities (Kawana et al., 2017). Similarly, among individual cardiomyocytes, R723G showed heterogeneity in cell-cell interaction and force generation, presumably due to differential levels of myosin proteins in each cell (Kraft et al., 2016). Surprisingly, we identified a cascade of fibrotic transcripts to be down-regulated in the mutant cardiomyocytes. Since fibrosis and myocyte disarray are hallmark features of HCM (Maron and Maron, 2013; Varnava et al., 2000), we believe that the down-regulation of fibrotic pathways could be a compensatory mechanism to reduce the functional load on the mutant cardiomyocytes.

It should be noted that genetic testing for patients with an HCM phenotype is recommended in the current guidelines and assesses 8 sarcomeric genes with strong evidence to be disease causing in HCM, including *MYH7* (Ommen et al., 2020). Genetic testing can be utilized for screening family members who may be at risk for developing HCM, however mutation type is not currently able to predict the risk of SCD or guide clinical management. However, genetic testing can be clinically important in those patients with greater than 1 sarcomeric gene mutation and HCM phenotype, as they may have more severe disease (Girolami *et al*., 2010; Kelly and Semsarian, 2009; Maron *et al*., 2012; Weissler-Snir *et al*., 2017). Prior studies using hiPSC-CMs have focused on single sarcomeric HCM mutations (Han et al., 2014; Lan *et al*., 2013). Here we have evaluated the impact of two sarcomeric mutations in hiPSC-CM, including a known pathogenic mutation (*MYH7* R723C) and a variant of unknown significance (*MYH6* R725C). We have assessed the molecular and physiologic impact of the double and single pathogenic mutations at an early-stage including calcium dysregulation, cellular hypertrophy, altered energetics, and altered extracellular matrix dynamics. While *MYH6* mutations have been described to result in HCM, they are not routinely tested in the sarcomeric gene panel. Patients with other sarcomeric gene mutations may have other relevant HCM causing gene mutations that are not being captured with routine genetic testing. Our study provides evidence of later-stage HCM phenotype with dual mutations and suggests developing a broader panel of mutations or whole genome sequencing to screen individuals with a genetic propensity for HCM or with advanced symptoms may improve the ability of genetic testing to guide treatment and/or predict disease penetrance.

Because hiPSC-CMs are relatively immature when cultured in 2D without exogenous stimulation, they uniquely facilitate the understanding of early-stage, direct consequences of HCM mutation(s) without other secondary changes in animal models or patients. Based on single-cell 2D cardiac differentiation showing that *MYH6* is abundant at day 12 and subsequently *MYH7* predominates (Grancharova et al., 2021), we assessed an early time point (Day 15) and a later time point (Day 45) to facilitate the early-stage understanding of the functional contributions of *MYH6* as well as *MYH7* converter domain mutations. Additionally, generation of hiPSCs with multiple HCM mutations including variants of unknown significance or sarcomere gene negative HCM, may allow understanding of early stage molecular and physiologic changes that have been challenging to assess in animal models or patients. Further, modeling double or multiple HCM mutations in hiPSC-CMs may elucidate the interaction between multiple sarcomeric proteins.

In summary, our data demonstrated that HCM phenotypes can be recapitulated using hiPSC-CMs with *MYH7* R723C and *MYH6* R725C mutations, and that changes in gene expression at early stages of differentiation could offer clues for novel therapeutic classes to prevent or delay the onset of HCM cellular and functional phenotypes. Future work to generate *MYH7* R723C mutant models in a 3D environment, like the human chambered muscle pump (hChaMP) (Kupfer et al., 2020) or engineered heart tissue (Yang et al., 2018), will help to further elucidate the role of the ECM in driving the phenotypic characteristics of HCM.

## 5 Data Availability Statement

All datasets generated for this study are included in the article/Supplementary Material.

## 6 Author Contributions

J.H., K.B., E.R.K., B.R.W., B.N.S., B.M.O. conceived and designed the research. K.B., B.S.M., and B.R.W. generated the R723C mutant hiPSCs. J.H. designed and executed the experiments including cardiac differentiation, functional measurements, endpoint sample collection, immunostaining, imaging and quantitative analysis, RNAseq analysis and flow cytometry analysis. B.N.S. performed qPCR analysis, ATP/ADP assay, pFAK staining and interpreted RNAseq result. F.K. helped with the clinical data analysis and summarized the clinical outcomes. S.G. analyzed blinded data for multinucleation and sarcomeric organization. J.E.A.L. helped interpret RNAseq result. J.H., F.K., B.N.S. and B.M.O. discussed the results, wrote and edited the manuscript. All authors approved the manuscript.

## 8 Acknowledgements

We acknowledge the help from Wei-Han Lin for carrying out the contractility and calcium assays. We acknowledge Dr. Ryan Knoper for helpful discussion about the clinical manifestation and characteristics of hypertrophic cardiomyopathy.

## 9 Supplementary Material

The Supplementary Material for this article can be found online at:

*The authors declare that the research was conducted in the absence of any commercial or financial relationships that could be construed as a potential conflict of interest*.

## Figure Legends

**Video S1. This video shows a side-by-side comparison of morphology and beat profile in *MYH7/MYH6* mutant hiPSC-CMs and isogenic control hiPSC-CMs**.

## References

Alpert, N.R., Brosseau, C., Federico, A., Krenz, M., Robbins, J., and Warshaw, D.M. (2002). Molecular mechanics of mouse cardiac myosin isoforms. Am J Physiol Heart Circ Physiol 283, H1446–1454. 10.1152/ajpheart.00274.2002.

Becker, K.D., Gottshall, K.R., Hickey, R., Perriard, J.C., and Chien, K.R. (1997). Point mutations in human beta cardiac myosin heavy chain have differential effects on sarcomeric structure and assembly: an ATP binding site change disrupts both thick and thin filaments, whereas hypertrophic cardiomyopathy mutations display normal assembly. J Cell Biol 137, 131–140. 10.1083/jcb.137.1.131.

Bhagwan, J.R., Mosqueira, D., Chairez-Cantu, K., Mannhardt, I., Bodbin, S.E., Bakar, M., Smith, J.G.W., and Denning, C. (2020). Isogenic models of hypertrophic cardiomyopathy unveil differential phenotypes and mechanism-driven therapeutics. J Mol Cell Cardiol 145, 43–53. 10.1016/j.yjmcc.2020.06.003.

Burke, M.A., Cook, S.A., Seidman, J.G., and Seidman, C.E. (2016). Clinical and Mechanistic Insights Into the Genetics of Cardiomyopathy. J Am Coll Cardiol 68, 2871–2886. 10.1016/j.jacc.2016.08.079.

Gaudelli, N.M., Komor, A.C., Rees, H.A., Packer, M.S., Badran, A.H., Bryson, D.I., and Liu, D.R. (2017). Programmable base editing of A*T to G*C in genomic DNA without DNA cleavage. Nature 551, 464–471. 10.1038/nature24644.

Girolami, F., Ho, C.Y., Semsarian, C., Baldi, M., Will, M.L., Baldini, K., Torricelli, F., Yeates, L., Cecchi, F., Ackerman, M.J., and Olivotto, I. (2010). Clinical features and outcome of hypertrophic cardiomyopathy associated with triple sarcomere protein gene mutations. J Am Coll Cardiol 55, 1444–1453. 10.1016/j.jacc.2009.11.062.

Grancharova, T., Gerbin, K.A., Rosenberg, A.B., Roco, C.M., Arakaki, J.E., DeLizo, C.M., Dinh, S.Q., Donovan-Maiye, R.M., Hirano, M., Nelson, A.M., et al. (2021). A comprehensive analysis of gene expression changes in a high replicate and open-source dataset of differentiating hiPSC-derived cardiomyocytes. Sci Rep 11, 15845. 10.1038/s41598-021-94732-1.

Guan, J.L. (1997). Role of focal adhesion kinase in integrin signaling. Int J Biochem Cell Biol 29, 1085–1096. 10.1016/s1357-2725(97)00051-4.

Guan, J.L. (2010). Integrin signaling through FAK in the regulation of mammary stem cells and breast cancer. IUBMB Life 62, 268–276. 10.1002/iub.303.

Han, L., Li, Y., Tchao, J., Kaplan, A.D., Lin, B., Li, Y., Mich-Basso, J., Lis, A., Hassan, N., London, B., et al. (2014). Study familial hypertrophic cardiomyopathy using patient-specific induced pluripotent stem cells. Cardiovasc Res 104, 258–269. 10.1093/cvr/cvu205.

Homburger, J.R., Green, E.M., Caleshu, C., Sunitha, M.S., Taylor, R.E., Ruppel, K.M., Metpally, R.P., Colan, S.D., Michels, M., Day, S.M., et al. (2016). Multidimensional structure-function relationships in human beta-cardiac myosin from population-scale genetic variation. Proc Natl Acad Sci U S A 113, 6701–6706. 10.1073/pnas.1606950113.

Kawana, M., Sarkar, S.S., Sutton, S., Ruppel, K.M., and Spudich, J.A. (2017). Biophysical properties of human beta-cardiac myosin with converter mutations that cause hypertrophic cardiomyopathy. Sci Adv 3, e1601959. 10.1126/sciadv.1601959.

Kelly, M., and Semsarian, C. (2009). Multiple mutations in genetic cardiovascular disease: a marker of disease severity? Circ Cardiovasc Genet 2, 182–190. 10.1161/CIRCGENETICS.108.836478.

Kohler, J., Winkler, G., Schulte, I., Scholz, T., McKenna, W., Brenner, B., and Kraft, T. (2002). Mutation of the myosin converter domain alters cross-bridge elasticity. Proc Natl Acad Sci U S A 99, 3557–3562. 10.1073/pnas.062415899.

Kraft, T., Montag, J., Radocaj, A., and Brenner, B. (2016). Hypertrophic Cardiomyopathy: Cell-to-Cell Imbalance in Gene Expression and Contraction Force as Trigger for Disease Phenotype Development. Circ Res 119, 992–995. 10.1161/CIRCRESAHA.116.309804.

Kupfer, M.E., Lin, W.H., Ravikumar, V., Qiu, K., Wang, L., Gao, L., Bhuiyan, D., Lenz, M., Ai, J., Mahutga, R.R., et al. (2020). In Situ Expansion, Differentiation and Electromechanical Coupling of Human Cardiac Muscle in a 3D Bioprinted, Chambered Organoid. Circ Res. 10.1161/CIRCRESAHA.119.316155.

Lan, F., Lee, A.S., Liang, P., Sanchez-Freire, V., Nguyen, P.K., Wang, L., Han, L., Yen, M., Wang, Y., Sun, N., et al. (2013). Abnormal calcium handling properties underlie familial hypertrophic cardiomyopathy pathology in patient-specific induced pluripotent stem cells. Cell Stem Cell 12, 101–113. 10.1016/j.stem.2012.10.010.

Lian, X., Hsiao, C., Wilson, G., Zhu, K., Hazeltine, L.B., Azarin, S.M., Raval, K.K., Zhang, J., Kamp, T.J., and Palecek, S.P. (2012). Robust cardiomyocyte differentiation from human pluripotent stem cells via temporal modulation of canonical Wnt signaling. Proc Natl Acad Sci U S A 109, E1848–1857. 10.1073/pnas.1200250109.

Locher, M.R., Razumova, M.V., Stelzer, J.E., Norman, H.S., Patel, J.R., and Moss, R.L. (2009). Determination of rate constants for turnover of myosin isoforms in rat myocardium: implications for in vivo contractile kinetics. Am J Physiol Heart Circ Physiol 297, H247–256. 10.1152/ajpheart.00922.2008.

Lowes, B.D., Minobe, W., Abraham, W.T., Rizeq, M.N., Bohlmeyer, T.J., Quaife, R.A., Roden, R.L., Dutcher, D.L., Robertson, A.D., Voelkel, N.F., et al. (1997). Changes in gene expression in the intact human heart. Downregulation of alpha-myosin heavy chain in hypertrophied, failing ventricular myocardium. J Clin Invest 100, 2315–2324. 10.1172/JCI119770.

Lowey, S., Lesko, L.M., Rovner, A.S., Hodges, A.R., White, S.L., Low, R.B., Rincon, M., Gulick, J., and Robbins, J. (2008). Functional effects of the hypertrophic cardiomyopathy R403Q mutation are different in an alpha-or beta-myosin heavy chain backbone. J Biol Chem 283, 20579–20589. 10.1074/jbc.M800554200.

Marian, A.J., and Braunwald, E. (2017). Hypertrophic Cardiomyopathy: Genetics, Pathogenesis, Clinical Manifestations, Diagnosis, and Therapy. Circ Res 121, 749–770. 10.1161/CIRCRESAHA.117.311059.

Maron, B.J., and Maron, M.S. (2013). Hypertrophic cardiomyopathy. Lancet 381, 242-255. 10.1016/S0140-6736(12)60397-3.

Maron, B.J., Maron, M.S., and Semsarian, C. (2012). Double or compound sarcomere mutations in hypertrophic cardiomyopathy: a potential link to sudden death in the absence of conventional risk factors. Heart Rhythm 9, 57–63. 10.1016/j.hrthm.2011.08.009.

McNally, E.M., Gianola, K.M., and Leinwand, L.A. (1989). Complete nucleotide sequence of full length cDNA for rat alpha cardiac myosin heavy chain. Nucleic Acids Res 17, 7527–7528. 10.1093/nar/17.18.7527.

Mosqueira, D., Mannhardt, I., Bhagwan, J.R., Lis-Slimak, K., Katili, P., Scott, E., Hassan, M., Prondzynski, M., Harmer, S.C., Tinker, A., et al. (2018). CRISPR/Cas9 editing in human pluripotent stem cell-cardiomyocytes highlights arrhythmias, hypocontractility, and energy depletion as potential therapeutic targets for hypertrophic cardiomyopathy. Eur. Heart J. 39, 3879–3892. 10.1093/eurheartj/ehy249.

Ommen, S.R., Mital, S., Burke, M.A., Day, S.M., Deswal, A., Elliott, P., Evanovich, L.L., Hung, J., Joglar, J.A., Kantor, P., et al. (2020). 2020 AHA/ACC Guideline for the Diagnosis and Treatment of Patients With Hypertrophic Cardiomyopathy: A Report of the American College of Cardiology/American Heart Association Joint Committee on Clinical Practice Guidelines. J Am Coll Cardiol 76, e159–e240. 10.1016/j.jacc.2020.08.045.

Rose, J., Kraft, T., Brenner, B., and Montag, J. (2020). Hypertrophic cardiomyopathy MYH7 mutation R723G alters mRNA secondary structure. Physiol Genomics 52, 15–19. 10.1152/physiolgenomics.00100.2019.

Sedaghat-Hamedani, F., Kayvanpour, E., Tugrul, O.F., Lai, A., Amr, A., Haas, J., Proctor, T., Ehlermann, P., Jensen, K., Katus, H.A., and Meder, B. (2018). Clinical outcomes associated with sarcomere mutations in hypertrophic cardiomyopathy: a meta-analysis on 7675 individuals. Clin Res Cardiol 107, 30–41. 10.1007/s00392-017-1155-5.

Semsarian, C., Ingles, J., Maron, M.S., and Maron, B.J. (2015). New perspectives on the prevalence of hypertrophic cardiomyopathy. J Am Coll Cardiol 65, 1249–1254. 10.1016/j.jacc.2015.01.019.

Sewanan, L.R., Schwan, J., Kluger, J., Park, J., Jacoby, D.L., Qyang, Y., and Campbell, S.G. (2019). Extracellular Matrix From Hypertrophic Myocardium Provokes Impaired Twitch Dynamics in Healthy Cardiomyocytes. JACC Basic Transl Sci 4, 495–505. 10.1016/j.jacbts.2019.03.004.

Varnava, A.M., Elliott, P.M., Sharma, S., McKenna, W.J., and Davies, M.J. (2000). Hypertrophic cardiomyopathy: the interrelation of disarray, fibrosis, and small vessel disease. Heart 84, 476–482. 10.1136/heart.84.5.476.

Webber, B.R., Lonetree, C.L., Kluesner, M.G., Johnson, M.J., Pomeroy, E.J., Diers, M.D., Lahr, W.S., Draper, G.M., Slipek, N.J., Smeester, B.A., et al. (2019). Highly efficient multiplex human T cell engineering without double-strand breaks using Cas9 base editors. Nat Commun 10, 5222. 10.1038/s41467-019-13007-6.

Weissler-Snir, A., Adler, A., Williams, L., Gruner, C., and Rakowski, H. (2017). Prevention of sudden death in hypertrophic cardiomyopathy: bridging the gaps in knowledge. Eur Heart J 38, 1728–1737. 10.1093/eurheartj/ehw268.

Yang, K.C., Breitbart, A., De Lange, W.J., Hofsteen, P., Futakuchi-Tsuchida, A., Xu, J., Schopf, C., Razumova, M.V., Jiao, A., Boucek, R., et al. (2018). Novel Adult-Onset Systolic Cardiomyopathy Due to MYH7 E848G Mutation in Patient-Derived Induced Pluripotent Stem Cells. JACC Basic Transl Sci 3, 728–740. 10.1016/j.jacbts.2018.08.008.

