## Supplementary Material for "Myosin Heavy Chain Converter Domain Mutations Drive Early-Stage Changes in Extracellular Matrix Dynamics in Hypertrophic Cardiomyopathy"

#### 1 Supplementary Figures

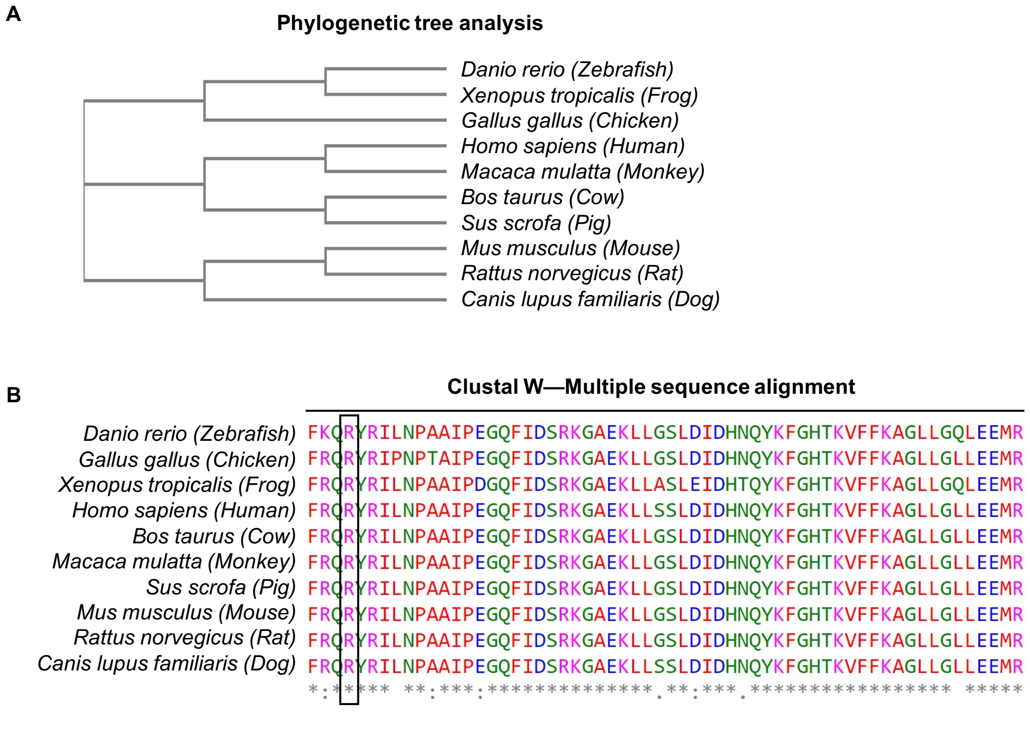


**Figure S1. Species comparison of *MYH7***

**(A)** Phylogenetic tree analysis showing divergence of the human and mouse gene and **(B)** multiple-alignment using Clustal W showing the conserved Arginine residue at 723 position across different species.

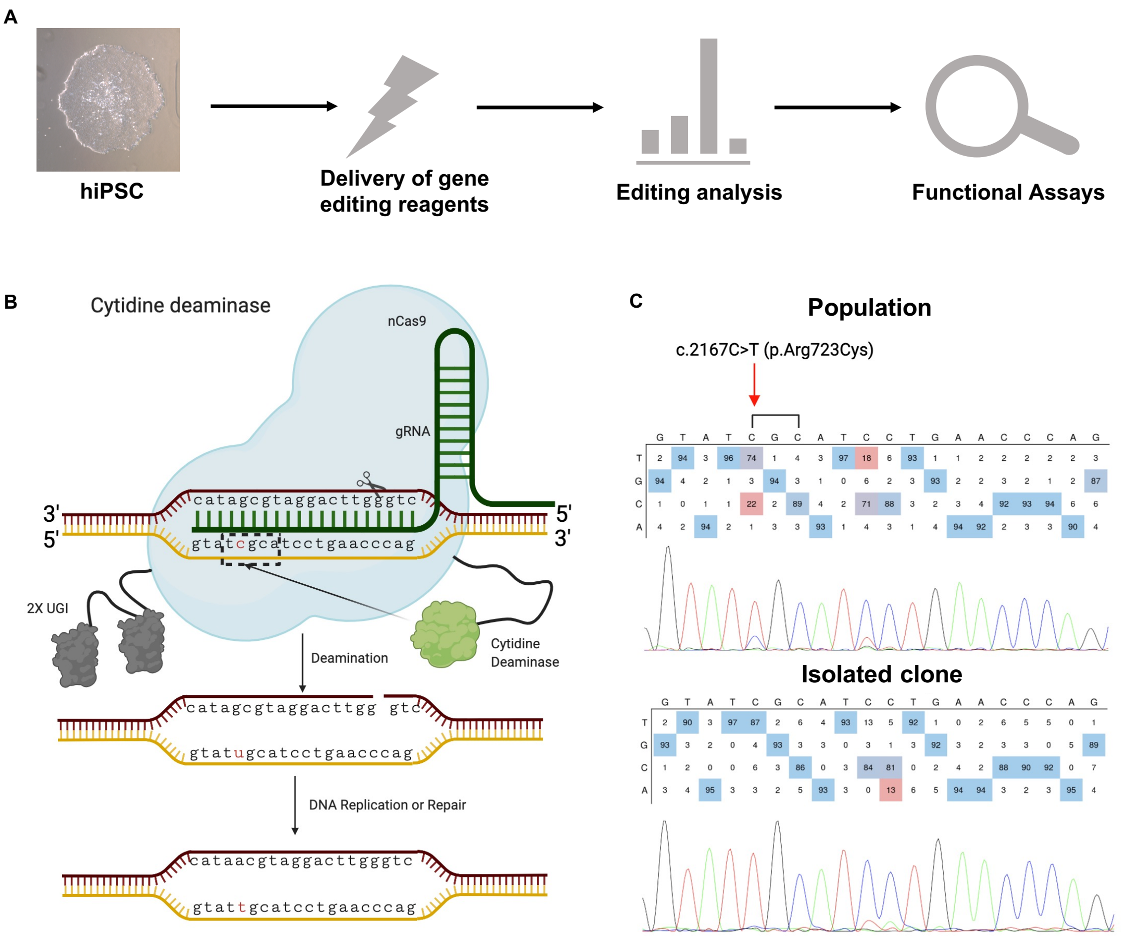


#### Figure S2. Base editing to create the *MYH7* c.2167C>T conversion without double strand breaks

**(A)** Schematic of workflow for creating the *MYH7* c.2167C>T conversion in hiPSCs. **(B)** Base editing process using cytidine base editor (CBE) with a target cytidine (c) (red). Briefly, nuclease Cas9 (blue) was directed to the *MYH7* c.2167C>T conversion site by the sgRNA (green) complementary to the target strand (gtatcgcatcctgaacccag). A cytidine deaminase converts the target c to uracil (u) which is read by DNA repair enzymes as the correct base due to the nick in the non-target strand caused by Cas9, resulting in u conversion to thymine (t). **(C)** Base editing results, with nucleotide ratios (top) and chromatogram (bottom). In the bulk population 74% of the target c were successfully converted to t. Clones were then isolated from this bulk population. Note that we followed the same process to create the *MYH6* c.2173C>T conversion and isolated *MYH7/MYH6* mutant clone via clonal selection.

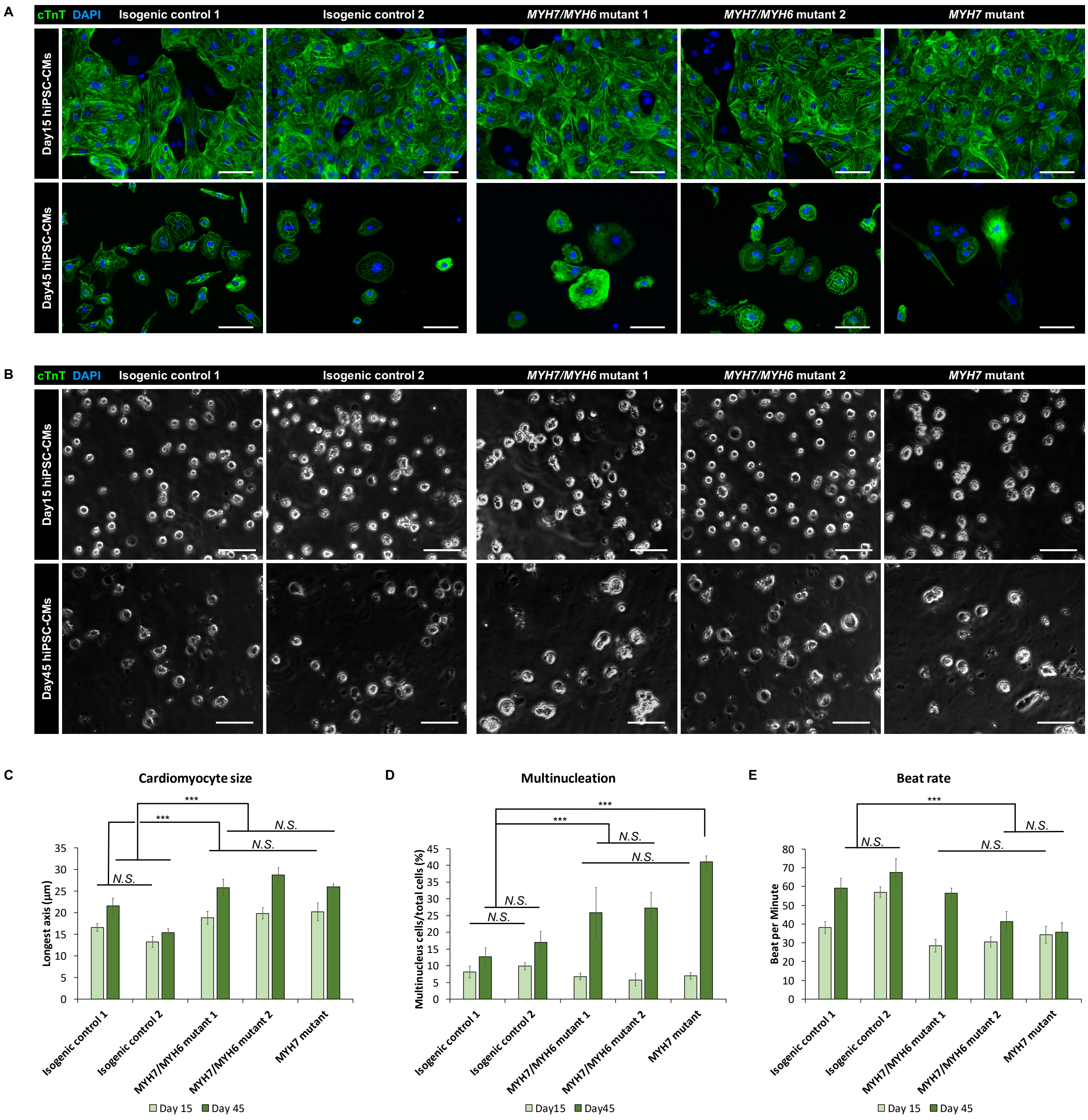


### Figure S3 HCM phenotypes in multiple isogenic control and mutant hiPSC-CMs

**(A)** Representative images of cTnT and DAPI-stained hiPSC-CMs on Day 15 and 45 of differentiation. Isogenic control 1 and *MYH7/MYH6* mutant 2 are the original lines shown in the main text. Isogenic control 2 shared the same parental hiPSC line with isogenic control 1, while isogenic control 2 does not over express CCND2. *MYH7/MYH6* mutant 1 has both *MYH7* R723C and *MYH6* R725C mutations as *MYH7/MYH6* mutant 2, the only difference between *MYH7/MYH6* mutant 1 and 2 is that they are two different isolated clones after base editing. *MYH7* mutant line only has *MYH7* R723C mutation, thus the difference between *MYH7* mutant and *MYH7/MYH6* mutant could indicate the impact of *MYH6* R725C. (Scale Bar = 100 μm) **(B)** Bright-field images of the re-plated hiPSC-CMs prior to their attachment to the substrate to analyze cardiomyocyte size at Day 15 and 45 of differentiation. (Scale Bar = 100 μm) **(C)** Quantification of cardiomyocyte size. Measured 24 cells that were randomly picked from the bright-field images for each condition. Note that no significant difference between isogenic control 1 and 2 at Day 15 of differentiation, and no significant difference among the *MYH7/MYH6* mutant 1, *MYH7/MYH6* mutant 2 and *MYH7* mutant. Statistically significant differences between any of the isogenic control and any of the mutant at both Day 15 and 45 time points. **(D)** Quantification of multinucleation. Analyzed from 9 IHC images for each condition. Note that no significant difference between isogenic control 1 and 2 at Day 15 and 45 of differentiation, no significant difference among the *MYH7/MYH6* mutant 1, *MYH7/MYH6* mutant 2 and *MYH7* mutant at Day 15, and no significant difference between *MYH7/MYH6* mutant 1 and 2 at Day 45. Statistically significant differences between any of the isogenic control and any of the mutant at Day 45 of differentiation. **(E)** Quantification of beat rate. Analyzed from 18 calcium transient measurements for each condition. Note that no significant difference between isogenic control 1 and 2 at Day 45 of differentiation, no significant difference among the *MYH7/MYH6* mutant 1, *MYH7/MYH6* mutant 2 and *MYH7* mutant at Day 15, and no significant difference between *MYH7/MYH6* mutant 2 and *MYH7* mutant at Day 45. Statistically significant differences between any of the isogenic control and *MYH7/MYH6* mutant 2 or *MYH7* mutant at Day 45 of differentiation. Data represents mean ± SEM. (*N.S.*: not significant; ****P*<0.005).

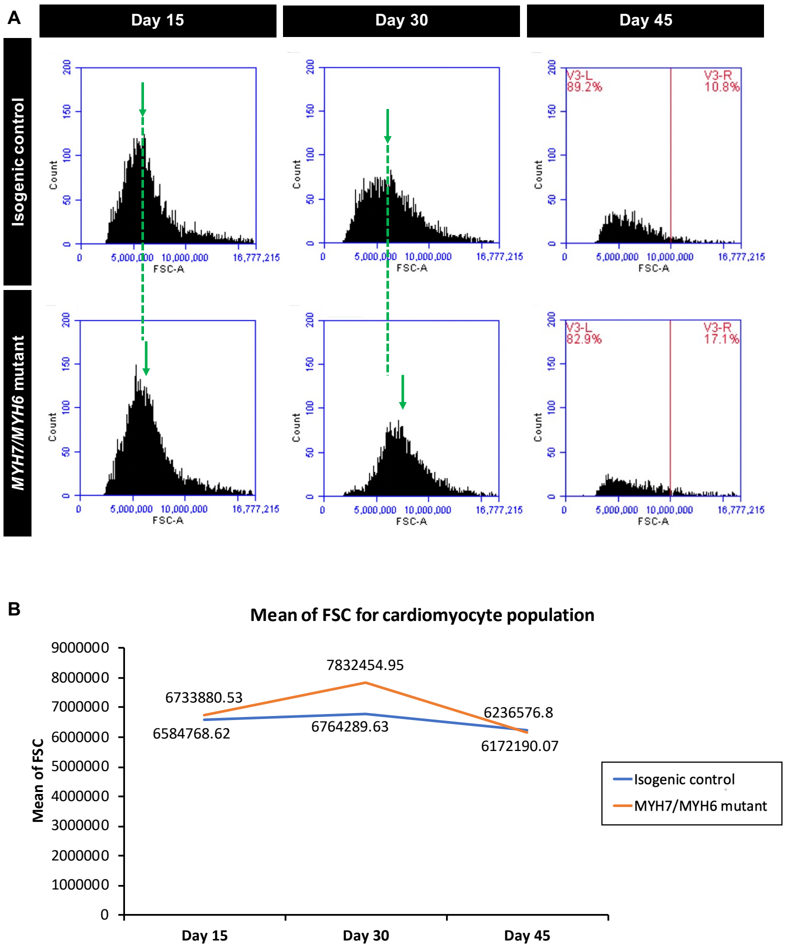

**Figure S4. FACS analysis for cardiomyocyte size**

**(A)** Event count for forward scatter (FSC) for both isogenic control and *MYH7/MYH6* mutant hiPSC-CMs on Day 15, Day 30 and Day 45 time points. Green arrows and dash lines indicate peak location. Red lines were arbitrarily assigned to locate at FSC = 10,000,000 for showing the significantly higher percentage of larger cell population in the *MYH7/MYH6* mutant hiPSC-CMs. **(B)** Mean scatter over time; higher values indicate larger cells. The *MYH7/MYH6* mutant hiPSC-CMs are larger at Day 30 than the isogenic control.

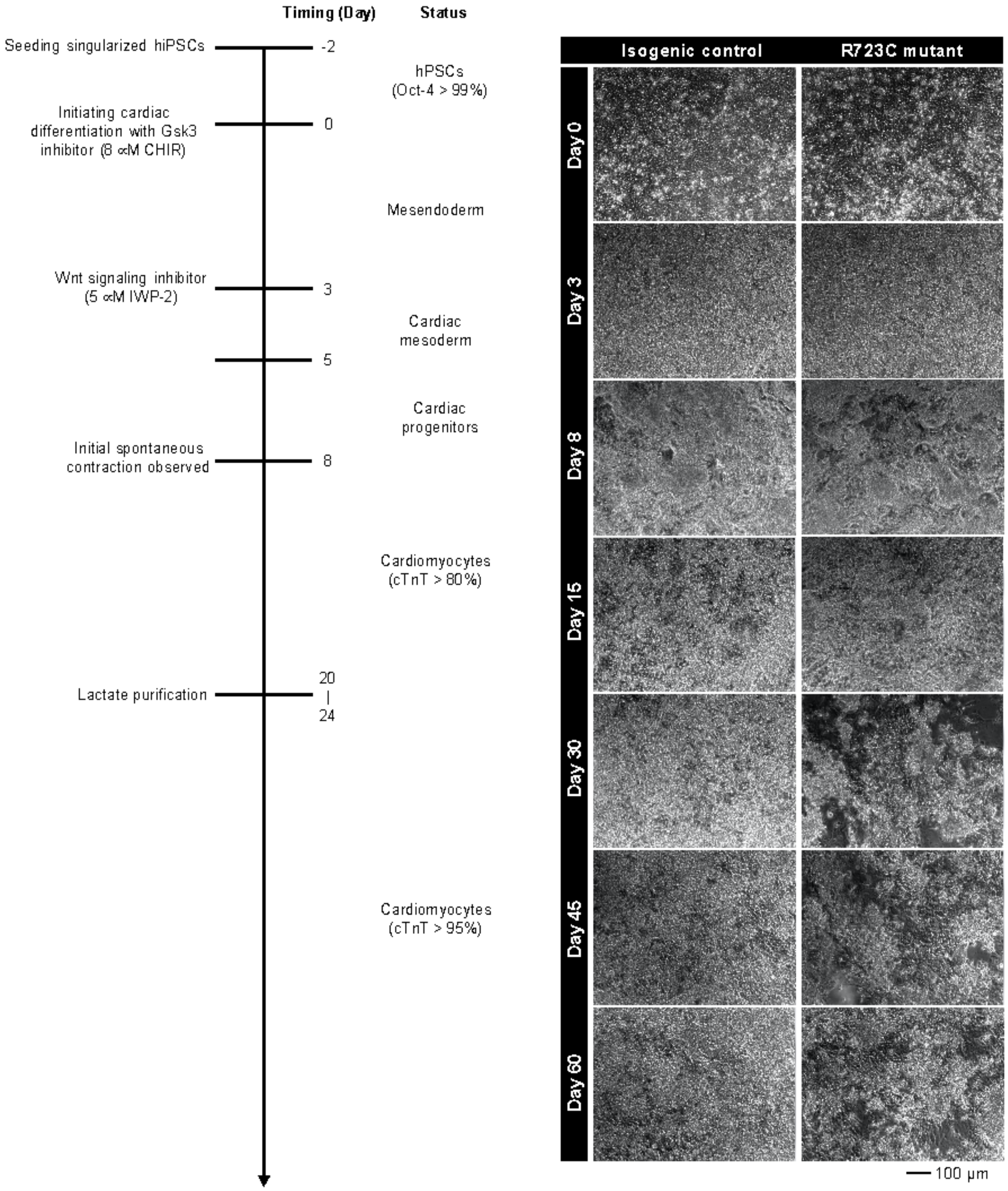


**Figure S5. Cardiac differentiation protocol with bright field images on multiple time points**

Supplementary Material

Bright field images of the cell morphology for both isogenic control and *MYH7/MYH6* mutant lines on Day 0, Day 3, Day 8, Day 15, Day 30, Day 45 and Day 60 after the start of differentiation. (Bar = 100 μm)

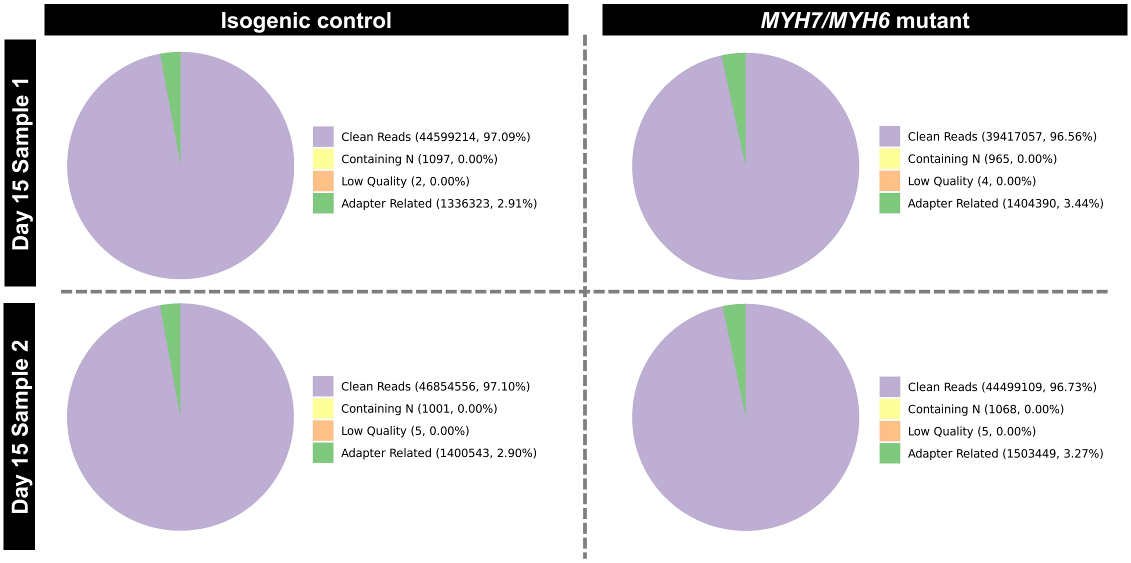


**Figure S6. RNASeq quality control**

Similarity of the read counts between isogenic control and mutant lines at Day 15 of differentiation.

A

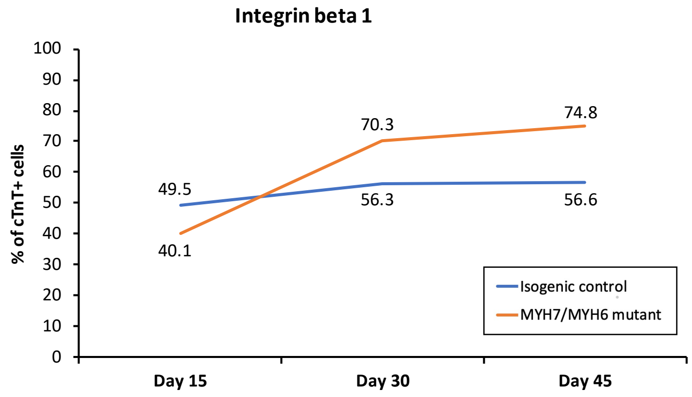


| Day 15 RNAseq (mutant vs control) | |
|---|---|
| *ITGB1* (ENSG00000150093) | |
| $log_2$(Fold Change) | *p*-value |
| 0.141 | $2.40 \times 10^{-02}$ |

B

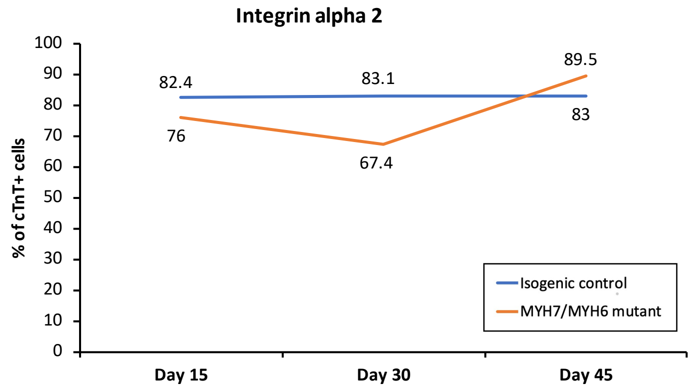


| Day 15 RNAseq (mutant vs control) | |
|---|---|
| *ITGA2* (ENSG00000164171) | |
| $log_2$(Fold Change) | *p*-value |
| -1.380 | $2.46 \times 10^{-10}$ |

C

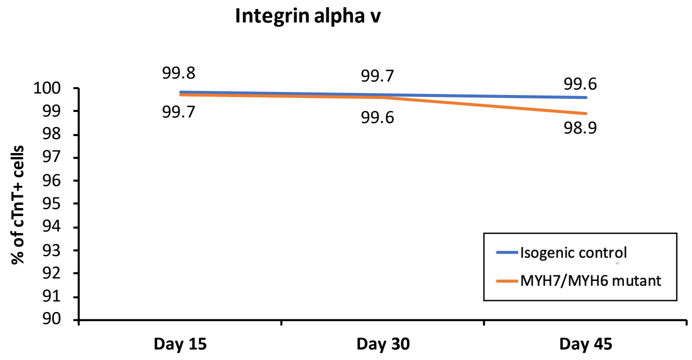


| Day 15 RNAseq (mutant vs control) | |
|---|---|
| *ITGAV* (ENSG00000138448) | |
| $log_2$(Fold Change) | *p*-value |
| 0.199 | $5.44 \times 10^{-03}$ |

**Figure S7. FACS analysis for integrin expression**

**(A)** Percentage of hiPSC-CMs expressing integrin beta 1 for both isogenic control and *MYH7/MYH6* mutant hiPSC-CMs on Day 15, Day 30 and Day 45 time points. **(B)** Percentage of hiPSC-CMs expressing integrin alpha 2 for both isogenic control and *MYH7/MYH6* mutant hiPSC-CMs on Day 15, Day 30 and Day 45 time points. **(C)** Percentage of hiPSC-CMs expressing integrin alpha v for both isogenic control and *MYH7/MYH6* mutant hiPSC-CMs on Day 15, Day 30 and Day 45 time points.

#### 2 Supplementary Tables

##### Table S1. Pathogenic Converter Domain Mutations and Associated hiPSC and Animal Studies

Mutations in MYH7 converter domain [residues 707-774] interpreted as "pathogenic" or "pathogenic/likely pathogenic" in the NCBI ClinVar database, as of April 2021.

| Table S1. Pathogenic Converter Domain Mutations and Associated hiPSC and Animal Studies | | | | |
|---|---|---|---|---|
| # | Mutation | Model | Alleles | Major Outcomes |
| 1 | **R723C** | Patient-derived hiPSC-CMs | Heterozygous | Telomere shortening in hiPSC-CMs (Chang et al., 2018) |
| 2 | **R723G** | hiPSC-CMs, swine model | Heterozygous | Hypertrophy, cell-cell imbalance, low survival in gene-edited pigs (Kraft et al., 2016; Montag et al., 2018) |
| 3 | **R719Q** | Patient-derived hiPSC-CMs | Heterozygous | Difference in structure and beating with non-isogenic control (Filippo Buono et al., 2020) |
| 4 | **R719W** | Patient-derived hiPSC-CMs, mouse MYH6 mutation | Heterozygous | Myosin conformation changes alter function, energetics and structure (Teekakirikul et al., 2010) |
| 5 | **G741R** | Mouse myoblasts | Heterozygous | Mutations did not affect myofibrillogenesis in myotubes (Miller et al., 2000) |
| 6 | **G716R** | Patient-derived hiPSC-CMs | Heterozygous | Telomere shortening in hiPSC-CMs (Chang et al., 2018) |

**Table S2. Clinical Characteristics of Patients with *MYH7* R723C mutation**

| Table S2. Clinical Characteristics of Patients with *MYH7* R723C mutation | | | | | | |
|---|---|---|---|---|---|---|
| # | Age | Sex | Pro-band | Symptoms and Outcomes | Cardiac Testing | Ref |
| 1* | 12 | F | N | **Symptoms:** Y<br>**Gene Penetrance:** Y<br>**Description:** End stage HF, Cardiac transplant | **MWT (mm):** 20<br>**LVED (mm):** 30 | 18 |
| 2 | 24 | M | Y | **Symptoms:** Y<br>**Gene Penetrance:** Y<br>**Description:** HF | **MWT (mm):** 38<br>**LVED (mm):** 40 | 17 |
| 3 | 40 | F | Y | **Symptoms:** Y<br>**Gene Penetrance:** Y<br>**Description:** Chest pain, Dyspnea, Syncope | **MWT (mm):** 38<br>**Echo result:** LVH | 6 |
| 4† | 39 | F | Y | **Symptoms:** N<br>**Gene Penetrance:** Y | **MWT (mm):** 22<br>**ECG result:** Lateral Q waves | 5 |
| 5† | 37 | F | N | **Symptoms:** N<br>**Gene Penetrance:** Y | **MWT (mm):** 27<br>**ECG result:** Inferior Q waves, TWI v1-v3, + SAM | 5 |
| 6† | 42 | M | N | **Symptoms:** N<br>**Gene Penetrance:** Y | **MWT (mm):** 22 | 5 |
| 7† | 43 | M | N | **Symptoms:** N<br>**Gene Penetrance:** Y | **MWT (mm):** 27<br>**ECG result:** Abnormal QRS axis | 5 |
| 8†ǂ | 9 | M | N | **Symptoms:** N<br>**Gene Penetrance:** Y | **MWT (mm):** 11<br>**ECG result:** Increased QRS voltage | 5 |
| 9†ǂ | 9 | M | N | **Symptoms:** N<br>**Gene Penetrance:** Not determined | **MWT (mm):** 8<br>**ECG result:** Increased QRS voltage | 5 |
| 10† | 17 | M | N | **Symptoms:** N<br>**Gene Penetrance:** Y | **MWT (mm):** 11 | 5 |
| 11† | 45 | F | N | **Symptoms:** N<br>**Gene Penetrance:** N | **MWT (mm):** 12.5 | 5 |
| 12† | 27 | F | N | **Symptoms:** N<br>**Gene Penetrance:** N | **MWT (mm):** 10 | 5 |
| 13† | 9 | M | N | **Symptoms:** N<br>**Gene Penetrance:** N | **MWT (mm):** 8 | 5 |
| 14† | 9 | M | N | **Symptoms:** N<br>**Gene Penetrance:** N | **MWT (mm):** 8 | 5 |

MWT = maximal wall thickness; SAM = systolic anterior motion; TWI = T-wave inversion; HF = heart failure; LVH = left ventricular hypertrophy; * father with same mutation and end stage HCM; † family; ǂ monozygotic twins.

Supplementary Material

**Table S3. List of primers used for qPCR analysis**

| # | Gene name (Accession number) | Primers (5'-3') | Tm | Amplicon length | Exon |
|---|---|---|---|---|---|
| 1 | **Human GAPDH** (NM_001256799.3) | Fwd- GGTCTCCTCTGACTTCAACAGCG<br>Rev- CCCTGTTGCTGTAGCCAAATTCG | 62<br>61 | 128 | 7-8 |
| 2 | **Human OCT4** (POU5F1) | Fwd- CCTGAAGCAGAAGAGGATCACCC<br>Rev- AAAGCGGCAGATGGTCGTTTG | 61<br>61 | 106 | 2-3 |
| 3 | **Human MYH7** (NM_000257.4) | Fwd- GGAGTTCACACGCCTCAAAGAGG<br>Rev- TCCTCAGCATCTGCCAGGTTGT | 61<br>63 | 147 | 22-23 |
| 4 | **Human MYH6** (NM_002471.3) | Fwd- GGAAGACAAGGTCAACAGCCTGT<br>Rev- TCCAGTTTCCGCTTTGCTCGC | 61<br>63 | 129 | 23-24 |
| 5 | **Human TNNI3** (NM_000363.5) | Fwd- CGTGTGGACAAGGTGGATGAAG<br>Rev- GCCGCTTAAACTTGCCTCGAAG | 61<br>60 | 118 | 5-6 |
| 6 | **Human ITGAV** (NM_002210.5) | Fwd- AGGAGAAGGTGCCTACGAAGCT<br>Rev- GCACAGGAAAGTCTTGCTAAGGC | 62<br>61 | 104 | 20-21 |
| 7 | **Human ITGA7** (NM_002206.3) | Fwd- CCTGTCCAATGAGAATGCCTCC<br>Rev- TCTACCTCCAGTTCCGTGGTCT | 60<br>60 | 132 | 16-17 |
| 8 | **Human ITGB1** (NM_002211.4) | Fwd- GGATTCTCCAGAAGGTGGTTTCG<br>Rev- TGCCACCAAGTTTCCCATCTCC | 60<br>61 | 143 | 6-7 |
| 9 | **Human ATP2B4** (NM_001684.5) | Fwd- CTCACCGAACTGACCTGTATCG<br>Rev- GGCTGTGTTGATGTTGTCACCTG | 60<br>61 | 135 | 12-13 |

### 3 Supplementary Video

**Supplementary Video S1.** This video shows a side-by-side comparison of morphology and beat profile in *MYH7/MYH6* mutant hiPSC-CMs and isogenic control hiPSC-CMs.

### 4 Supplementary Experimental Procedures

#### 4.1 Identification of patients with MYH7 R723C mutation and their clinical symptoms

We searched the ClinVar database and the Clinical Genome Resource database (https://erepo.clinicalgenome.org/evrepo/ui/interpretation/7cad6d45-ed00-44b9-a31d-225ff15a0c10) for publications containing references to *MYH7* Arg723Cys. A total of 23 manuscripts were reviewed and 4 articles were selected (O'Mahony et al., 2016; Rowin et al., 2014; Tesson et al., 1998; Watkins et al., 1992). The inclusion criteria were a single mutation of *MYH7* R723C and patient level clinical data. Double or triple mutations were excluded.

#### 4.2 Re-plating hiPSC-CMs

To obtain monolayer hiPSC-CMs, a re-plating process was performed. Briefly, 1 mL of 0.25% Trypsin was added to each well and incubated at 37°C for 20 min. Singularized cells were obtained by pipetting with P1000 tips several times. The cell suspension was transferred to a conical tube containing RPMI20 (RPMI1640 with L-glutamine and 20% FBS) with two-fold cell suspension volume. The suspension was centrifuged at 200×g for 5 min to remove supernatant and hiPSC-CMs were seeded at the density of $9 \times 10^4$ cells/well on 12-well plates in RPMI/B27 medium with 10μM ROCK inhibitor. The re-plated hiPSC-CMs were cultured for 48 h and then fixed with 4% PFA for immunostaining.

#### 4.3 Identification of Differentiation Efficiency

Differentiation efficiency was identified by the percentage of cardiomyocytes over total cell number. Cardiomyocytes were recognized via positive cTnT staining and total cell number was based on DAPI positive nuclei in a 10× view under the microscope (ZEISS, Oberkochen, Germany). At least 5 randomly picked 10× images for both mutant hiPSCCMs and control hiPSC-CMs were utilized for the calculation.

#### 4.4 Determination of hiPSC-CM Size

The singularized hiPSC-CMs were obtained via the same trypsinization process for re-plating. The cardiomyocyte size of the singularized hiPSC-CMs was examined by averaging the longest axis of each particle over 6 bright field images for each cell line on each time point.

#### 4.5 Calcium Transient Measurements

Movement of Calcium ions was assessed with DMi8 fluorescence microscope (Leica, Wetzlar, Germany) and LAS X software. Cultured hiPSC-CMs were incubated with 5 μM Fluo-4 acetoxymethyl ester (Fluo-4 AM) in culture medium at 37°C for 30 min, and then culture medium was exchanged to Tyrode's salt solution for another 30 min incubation at 37°C. After that, hiPSC-CMs were moved onto the microscope stage and covered with a heating plate to maintain the temperature at 37°C. Fluo-4 AM intensity was recorded at the frame rate of 6.90 Hz with a 30 ms exposure time. The acquired data was processed by ImageJ to obtain a time trace of calcium signal and then analyzed in a custom-written Python script to extract maximal and minimal intensity and corresponding time points for each peak. Peak amplitude was determined by $F/F_0$ where $F = \langle F_{max} \rangle - \langle F_{min} \rangle$ and $F_0 = \langle F_{min} \rangle$. $\langle F_{max} \rangle$ and $\langle F_{min} \rangle$ represent the averaged maximal and minimal intensity for each peak, respectively.

##### 4.6 Determination of Relative Cardiomyocyte Size and Integrin Expression via Flow Cytometry

Singularized cell samples were collected and fixed in 4% PFA for 15 min and then washed and stored in PBS at 4°C. At least $0.5 \times 10^6$ cells per sample were split and assigned to unstained control, secondary antibody only, cTnT only, and cTnT and integrin co-stained samples (integrin beta 1 or alpha 2 or alpha v). Primary antibodies (mouse anti-human cTnT, rabbit anti-human integrin beta 1, rabbit anti-human integrin alpha 2, and rabbit anti-human integrin alpha v) and secondary antibodies (goat anti-mouse 488, and goat anti-rabbit 647) were diluted in 10 μg/mL digitonin 1:100 and 1:500, respectively. Cell samples were permeabilized and immunostained simultaneously (30 min at 4°C for primary antibody, washed with 0.05% (v/v) Tween 20 in PBS twice and then 30 min at 4°C for secondary antibody). Washed with 0.05% (v/v) Tween 20 in PBS twice before resuspended in 200 μL of PBS. Acquire data by Flow Cytometry (BD Accuri C6 Flow Cytometer). To determine cardiomyocyte size and integrin expression level, gated each cell line on each time point with unstained control, adjusted with secondary antibody only control to exclude false positive events and then targeted cTnT+ events as cardiomyocytes for analysis.

##### 4.7 GO and KEGG Enrichment Analysis of Differentially Expressed Genes

Gene Ontology (GO) enrichment analysis of differentially expressed genes was implemented by the clusterProfiler R package, in which gene length bias was corrected. GO terms with corrected P-value less than 0.05 were considered significantly enriched by differential expressed genes. We used the clusterProfiler R package to test the statistical enrichment of differential expression genes in KEGG pathways.

Supplementary Material

#### SUPPLEMENTAL REFERENCES


Chang, A.C.Y., Chang, A.C.H., Kirillova, A., Sasagawa, K., Su, W., Weber, G., Lin, J., Termglinchan, V., Karakikes, I., Seeger, T., *et al.* (2018). Telomere shortening is a hallmark of genetic cardiomyopathies. Proc Natl Acad Sci U S A *115*, 9276-9281.

Filippo Buono, M., von Boehmer, L., Strang, J., Hoerstrup, S.P., Emmert, M.Y., and Nugraha, B. (2020). Human Cardiac Organoids for Modeling Genetic Cardiomyopathy. Cells *9*.

Kraft, T., Montag, J., Radocaj, A., and Brenner, B. (2016). Hypertrophic Cardiomyopathy: Cell-to-Cell Imbalance in Gene Expression and Contraction Force as Trigger for Disease Phenotype Development. Circ Res *119*, 992-995.

Miller, G., Colegrave, M., and Peckham, M. (2000). N232S, G741R and D778G beta-cardiac myosin mutants, implicated in familial hypertrophic cardiomyopathy, do not disrupt myofibrillar organisation in cultured myotubes. FEBS Lett *486*, 325-327.

Montag, J., Petersen, B., Flogel, A.K., Becker, E., Lucas-Hahn, A., Cost, G.J., Muhlfeld, C., Kraft, T., Niemann, H., and Brenner, B. (2018). Successful knock-in of Hypertrophic Cardiomyopathy-mutation R723G into the MYH7 gene mimics HCM pathology in pigs. Sci Rep *8*, 4786.

O'Mahony, C., Jichi, F., Monserrat, L., Ortiz-Genga, M., Anastasakis, A., Rapezzi, C., Biagini, E., Gimeno, J.R., Limongelli, G., McKenna, W.J., *et al.* (2016). Inverted U-Shaped Relation Between the Risk of Sudden Cardiac Death and Maximal Left Ventricular Wall Thickness in Hypertrophic Cardiomyopathy. Circ Arrhythm Electrophysiol *9*.

Rowin, E.J., Maron, B.J., Kiernan, M.S., Casey, S.A., Feldman, D.S., Hryniewicz, K.M., Chan, R.H., Harris, K.M., Udelson, J.E., DeNofrio, D., *et al.* (2014). Advanced heart failure with preserved systolic function in nonobstructive hypertrophic cardiomyopathy: under-recognized subset of candidates for heart transplant. Circ Heart Fail *7*, 967-975.

Teekakirikul, P., Eminaga, S., Toka, O., Alcalai, R., Wang, L., Wakimoto, H., Nayor, M., Konno, T., Gorham, J.M., Wolf, C.M., *et al.* (2010). Cardiac fibrosis in mice with hypertrophic cardiomyopathy is mediated by non-myocyte proliferation and requires Tgf-beta. J Clin Invest *120*, 3520-3529.

Tesson, F., Richard, P., Charron, P., Mathieu, B., Cruaud, C., Carrier, L., Dubourg, O., Lautie, N., Desnos, M., Millaire, A., *et al.* (1998). Genotype-phenotype analysis in four families with mutations in beta-myosin heavy chain gene responsible for familial hypertrophic cardiomyopathy. Hum Mutat *12*, 385-392.

Watkins, H., Thierfelder, L., Hwang, D.S., McKenna, W., Seidman, J.G., and Seidman, C.E. (1992). Sporadic hypertrophic cardiomyopathy due to de novo myosin mutations. J Clin Invest *90*, 1666-1671.
